# p57^Kip2^ acts as a transcriptional corepressor to regulate intestinal stem cell fate and proliferation

**DOI:** 10.1101/2022.09.09.507138

**Authors:** Justine Creff, Ada Nowosad, Anne Prel, Anne Pizzoccaro, Marion Aguirrebengoa, Nicolas Duquesnes, Caroline Callot, Thomas Jungas, Christine Dozier, Arnaud Besson

## Abstract

p57^Kip2^ is a cyclin/CDK inhibitor and a negative regulator of cell proliferation. Remarkably, p57 is the only CDK inhibitor required for embryonic development and p57 knockout mice display multiple developmental anomalies, including intestinal shortening. Here, we report that p57 regulates intestinal stem cell (ISC) fate and proliferation in a CDK-independent manner during intestinal development. In absence of p57, proliferation in intestinal crypts is markedly increased and genetic labelling experiments revealed an amplification of transit amplifying cells and of Hopx^+^ ISCs, which are no longer quiescent. On the other hand, Lgr5^+^ crypt-base columnar (CBC) cells were unaffected. RNA-Seq analyses of Hopx^+^ ISCs show major changes in gene expression in absence of p57. We found that p57 binds to and inhibits the activity of Ascl2, a transcription factor critical for ISC specification and maintenance, by participating in the recruitment of a corepressor complex to Ascl2 target gene promoters. Thus, our data suggests that during intestinal development, p57 plays a key role in maintaining Hopx^+^ stem cell quiescence and repressing the ISC phenotype outside of the crypt bottom by inhibiting the transcription factor Ascl2 in a CDK-independent manner.

## Introduction

p57^Kip2^, encoded by the *CDKN1C* gene, is a cyclin/CDK inhibitor of the Cip/Kip family with p21^Cip1/Waf1^ and p27^Kip1^. *CDKN1C* is an imprinted gene and p57 is expressed only from the maternal allele. Through inhibition of CDKs, p57 negatively regulates the cell division cycle, acting as a putative tumor suppressor [1, 2]. Unlike other CDK inhibitors, p57 plays a critical role during embryonic development and the knockout of p57 in mice (p57^KO^) results in perinatal lethality caused by numerous developmental anomalies [3, 4]. Some phenotypes of p57^KO^ mice resemble features observed in Beckwith-Wiedemann Syndrome (BWS) [4, 5]. Indeed, *CDKN1C* epigenetic repression or mutations are the most frequent causes of BWS [6, 7]. Like other Cip/Kip proteins, evidence indicate that in addition to its CDK inhibitory function, p57 regulates various biological processes by associating with different protein partners [2, 8]. Generation of a knock-in mouse model expressing a mutant form of p57 (p57^CK-^) that cannot bind cyclin/CDK complexes has provided genetic evidence of CDK-independent functions of p57 during embryonic development and revealed that BWS is not entirely caused by the loss of CDK inhibition [5]. Importantly, while approximately 30-40% of p57^KO^ mice exhibit intestinal shortening or truncation [3, 9], this phenotype is not observed in p57^CK-^ mutant mice [5], indicating that intestinal development is regulated by p57 in a CDK-independent manner. The mechanism underlying this role of p57 remains unknown. p57 expression was reported in the intestinal epithelium except at the bottom of crypts, where its expression is repressed by the transcription factor Hes1 downstream of Notch receptor signaling [10]. Disruption of Notch1 and -2 results in derepression of p27 and p57 and in premature cell cycle exit and differentiation of intestinal progenitors [10]. More recently, p57 expression was detected in populations of quiescent Mex3a^High^ and Bmi1^High^ ISCs, but its mechanism of action has not been studied [11, 12].

The intestinal epithelium is organized in crypt/villus structures and is one of the most rapidly renewing tissue in the body, with a turnover of 5 to 7 days [13, 14]. Villi are finger-like protrusions covered by a single epithelial layer of post-mitotic, differentiated cells. They are surrounded by crypts, which are invaginations into the mesenchyme where ISCs reside. The rapid renewal of the epithelium is driven by Lgr5^+^ ISCs, known as crypt-base columnar cells (CBCs), that localize at the bottom of the crypts, interspersed between Paneth cells [15]. Intestinal development starts around embryonic day 9.5 (E9.5) in mice with a single-layered pseudostratified epithelium lining the luminal surface of the gut tube [16]. At E14.5, after elongation and thickening of the intestinal tube, major remodeling occurs, resulting in the formation of non-proliferative villi and proliferative intervillous regions by E16.5 [17]. Lgr5^+^ ISCs first appear at E13.5 and localize to intervillous regions by E16.5 [18, 19]. Crypts and villi with all intestinal epithelial cell types are fully matured by postnatal day 15 (P15).

Lgr5^+^ ISCs divide symmetrically approximately once per day, and as these cells are progressively excluded from the stem cell zone, they become committed progenitors (secretory or absorptive) called transit-amplifying (TA) cells, that rapidly divide several times before terminally differentiating in the various epithelial cell types lining the intestine [14, 15, 20]. During differentiation, cells migrate upward from the crypt to the base of villi where they are fully specialized. Once in villi, upward migration continues until cells shed in the intestinal lumen and die by anoïkis upon reaching the top of villi [13, 21].

Despite their importance in homeostasis, acute loss of Lgr5^+^ ISCs does not compromise epithelial integrity. Indeed, due to their rapid cycling, Lgr5^+^ ISCs are sensitive to chemicals, irradiation or other damages but are regenerated within a few days at the base of the crypt and the epithelium is completely restored. However, constant depletion of Lgr5^+^ ISCs abrogates intestinal epithelium renewal, indicating they are the obligatory precursors to all intestinal cell types [22]. While several studies have shown by genetic lineage tracing that Lgr5^+^ ISC regeneration can be effected by multiple mature cell types that can dedifferentiate and divide to give new CBCs in the crypt, others have identified rare populations of quiescent reserve stem cells that can restore the Lgr5^+^ ISC pool when needed (Reviewed in [14, 23, 24]. Whether these reserve ISCs are “professional” stem cells or merely committed progenitors that can reverse to a CBC phenotype by dedifferentiation upon crypt injury is still debated.

A number of studies traced this regenerative capacity to a dedicated reserve stem cell pool existing normally in a quiescent state but able to activate and proliferate in case of damage to restore Lgr5^+^ ISCs. These reserve ISCs are located near the +4 position and have been identified using different markers, including Bmi1^+^ [25-27], mTert^+^ [28], Hopx^+^ [29] or Sox9^High^ [30, 31] or a rare population of Clusterin^+^ cells [32]. Furthermore, Lgr5^+^ and reserve ISCs can restore one another, highlighting a high level of plasticity in the crypt [29]. However, the existence and identity of reserve ISCs remains debated since none of the markers described for reserve ISCs are fully specific of these cells and at least some Lgr5^+^ ISCs express these markers [33-35]. Indeed, while most Lgr5^+^ ISCs are rapidly cycling, around 20% are quiescent and these cells were named LRCs (Label-Retaining Cells) [36]. LRCs exhibit both Lgr5^+^ and reserve ISCs gene expression signatures as well as a secretory progenitor gene signature [36]. Consistent with this, another study identified a population of largely quiescent (Ki67^-^) Lgr5^Low^ cells located around position +4 whose gene expression signature resembles that described for LRCs [37]. More recently, a Mex3a^High^ population was described as a quiescent Lgr5^+^ ISC subpopulation expressing both CBC and +4 markers involved in epithelial regeneration [11]. Another mechanism put forth by other studies is that upon damage, committed secretory (Dll1^+^) and absorptive (Alpi^+^) progenitors can undergo dedifferentiation and act as stem cells to regenerate Lgr5^+^ ISCs and the intestinal epithelium [38, 39]. Importantly, this dedifferentiation mechanism involves the re-expression and activation of Ascl2, a transcription factor required for Lgr5^+^ ISC specification and maintenance [40-42]. Regardless of whether Lgr5^+^ ISC recovery after damage is mediated by dedifferentiation of committed progenitors or “professional” reserve stem cells, how these cells exit quiescence and are mobilized to allow regeneration remains largely unclear. Here, we investigated the role of the CDK inhibitor p57^Kip2^ in intestinal homeostasis during development. We find that p57 controls the maintenance of quiescence in a progenitor/stem cell population in a CDK independent manner. Loss of p57 expression causes increased proliferation in crypts due to amplification of Hopx^+^ ISCs and TA cells, while Lgr5^+^ ISCs are not affected. Additionally, we found that p57 binds to and inhibits Ascl2 transcriptional activity by participating in the recruitment of a transcriptional corepressor complex containing HDAC7 and mSin3a at Ascl2 target gene promoters.

## Results

### p57 plays a critical role in intestinal development in a CDK independent manner

Approximately 30-40% of p57^KO^ mice exhibit intestinal shortening or truncation at E18.5 (Figure 1A-C) [3, 5, 9]. Importantly, as reported previously [5], this phenotype was not observed in p57^CK-^ mice in which p57 cannot bind to and inhibit cyclin-CDK complexes, (Figure 1A-C and Table S1), indicating that p57 plays a critical role in intestinal development independently of CDK inhibition. Histological analysis of H&E stained intestine sections shows that compared to p57^+/+^ and p57^CK-^ mice, p57^KO^ intestines exhibit a strong disorganization of the intestinal epithelium (Figure 1D), especially in villi where nuclear morphology was consistent with cell death. This was confirmed by immunostaining against cleaved caspase 3 (Figure S1A). Quantification of proliferative cells by Ki67 immunostaining revealed that p57^KO^ intestines exhibit a marked increase of proliferation in intervillous domains compared to p57^+/+^ mice (Figure 1E, F). Normal proliferation levels in p57^CK-^ intestines indicate that the p57^CK-^ protein is still able to limit proliferation in the crypt and that increased proliferation in p57^KO^ is not due to loss of CDK inhibition by p57 (Figure 1E, F). These results were confirmed by labeling proliferating cells with PCNA (Figure S1B). As previously reported [10], p57 expression was only observed in villi and not at the bottom of crypts, where CBCs reside, and p57^CK-^ had a similar expression pattern (Figure 1G and S1B). The tissue alterations and increased proliferation observed in p57^KO^ mice may result from an expansion of the stem cell compartment. To test this hypothesis, ISC populations were monitored using reporter mice to genetically label these cells.

**Figure 1:**
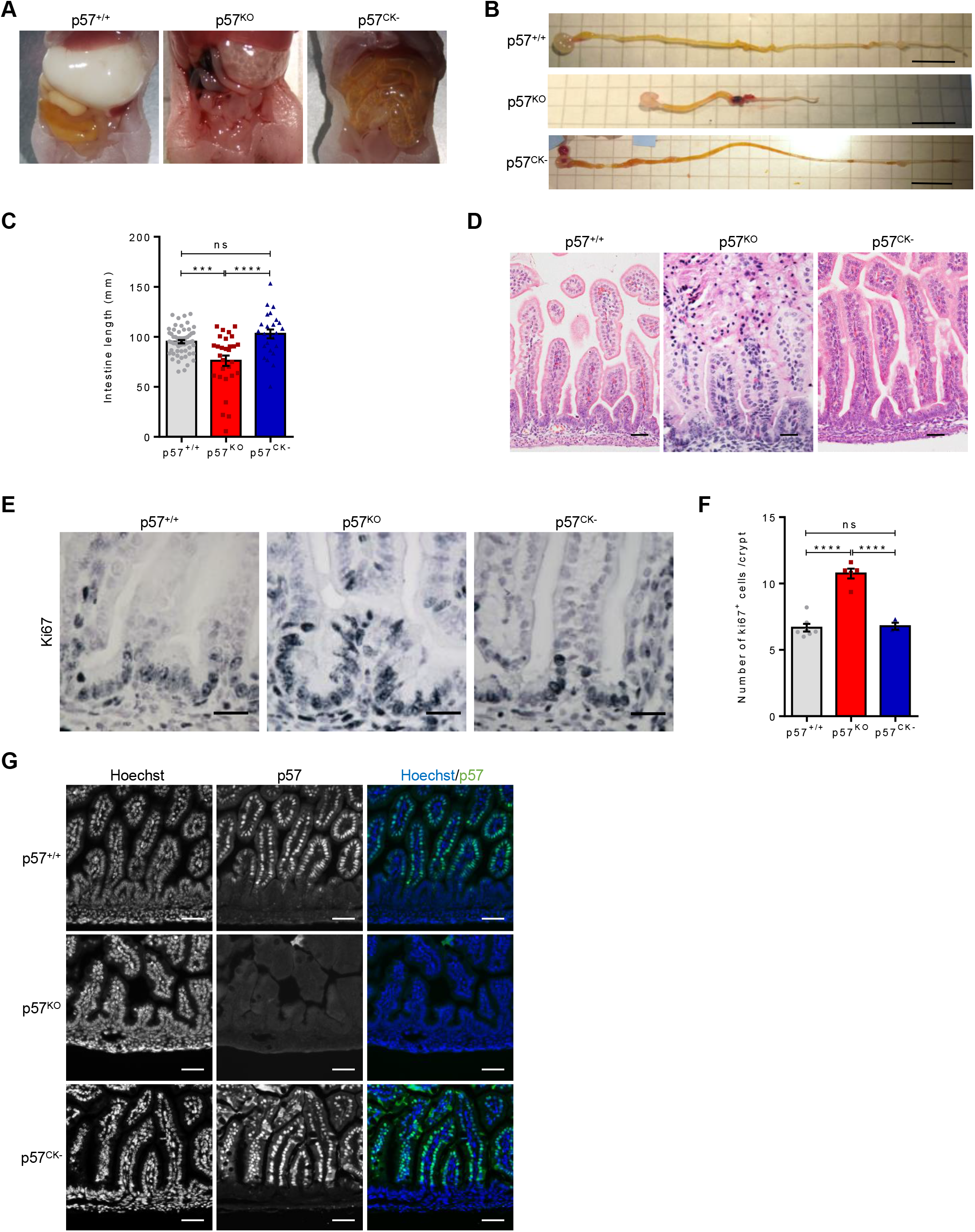
p57 plays a critical role in intestinal development in a CDK-independent manner. (A) Examples of images of abdomen of p57^+/+^, p57^KO^ and p57^CK-^ mice at P1. The presence of air in the gastrointestinal tract of p57^KO^ and p57^CK-^ mice is caused by cleft palate. (B) Examples of images of p57^+/+^, p57^KO^ and p57^CK-^ intestines at E18.5. Approximately 30% of p57^KO^ mice exhibit intestinal shortening, a phenotype never observed in p57^CK-^ mice. Scale bars, 1 cm. (C) Intestine length at E18.5 in p57^+/+^ (n=60), p57^KO^ (n=29) and p57^CK-^ (n=25) embryos. Graph shows means ± SEM. ns: p > 0.05; ***: p ≤ 0.001; ****: p ≤ 0.0001. (D) Representatives images of H&E stained intestine sections from E18.5 embryos showing intestinal epithelium alterations in p57^KO^ mice. (E) Representative images of Ki67 immunohistochemistry (grey) of intestine sections from p57^+/+^, p57^KO^ or p57^CK-^ E18.5 embryos. (F) Quantification of Ki67^+^ cells per intervillous domain from p57^+/+^ (n=7), p57^KO^ (n=5) and p57^CK-^ (n=3) E18.5 embryos as described in E. Graph shows means ± SEM. ns: p > 0.05; ****: p ≤ 0.0001. (G) Representative images of p57 immunostaining (green) of intestine sections from E18.5 p57^+/+^, p57^KO^ and p57^CK-^ embryos. DNA was stained with Hoechst 33342. Scale bars, 50 µm. See also Figure S1 and Table S1.

### p57 loss does not affect Lgr5^+^ ISCs

To label Lgr5^+^ ISCs, p57^KO^ and p57^CK-^ mice were intercrossed with Lgr5-eGFP-IRES-CreERT2 reporter mice (referred to as p57^+/+^;Lgr5^+/eGFP-CreERT2^, p57^KO^;Lgr5^+/eGFP-CreERT2^ and p57^CK-^;Lgr5^+/eGFP-CreERT2^), in which GFP expression is driven by the Lgr5 promoter, allowing detection of CBCs with GFP [15]. Evaluation of Lgr5^+^ ISC number by monitoring GFP on intestine cryosections did not reveal any significant difference between p57^+/+^, p57^CK-^ and p57^KO^ embryos (Figure 2A, B). Similar results were obtained by immunostaining for Olfm4 (Figure S2A, B), a Lgr5^+^ ISC marker [43]. By flow cytometry, this mouse model allows visualizing a GFP^High^ population that corresponds to CBCs and a GFP^Low^ population corresponding to their immediate progeny, the TA cells [33]. Consistent with the histological data, the percentage of GFP^High^ cells was not affected by p57 status (p57^+/+^: 0.15% ± 0.02; p57^KO^: 0.17% ± 0.04; p57^CK-^: 0.14% ± 0.01) (Figure 2C, D). On the other hand, the number of TA cells (GFP^Low^) was increased in p57^KO^ intestine (1.06% ± 0.18) compared to p57^+/+^ (0.58% ± 0.05) and p57^CK-^ (0.48% ± 0.13) animals (Figure 2C, D). These results were confirmed using Id1 immunostaining, which labels both ISCs and TA cells [44]. In agreement with the flow cytometry data, p57^KO^ intestine had more Id1^+^ cells than p57^+/+^ and p57^CK-^ mice (Figure 2E, F). The identity and molecular signature of Lgr5 GFP^High^, GFP^Low^ and GFP^Neg^ cells observed by flow cytometry was verified by RT-qPCR on sorted populations (Figure S2C-E). As expected, GFP^High^ cells exhibited strong expression of CBC markers such as Lgr5, Ascl2 and Olfm4, and low expression of differentiation markers (Muc2, Muc4 and Sucrase) (Figure S2C-D). These cells also expressed moderate levels of reserve ISC markers such as mTert, Sox9 and, to a lower extent, of Hopx and Mex3a (Figure S2E), as previously described [33, 34, 36]. Taken together, our data indicate that p57 does not affect Lgr5^+^ ISC abundance, consistent with p57 not being expressed in the lower crypt, however p57 loss causes an increase in the number of progenitor cells in a CDK-independent manner.

**Figure 2:**
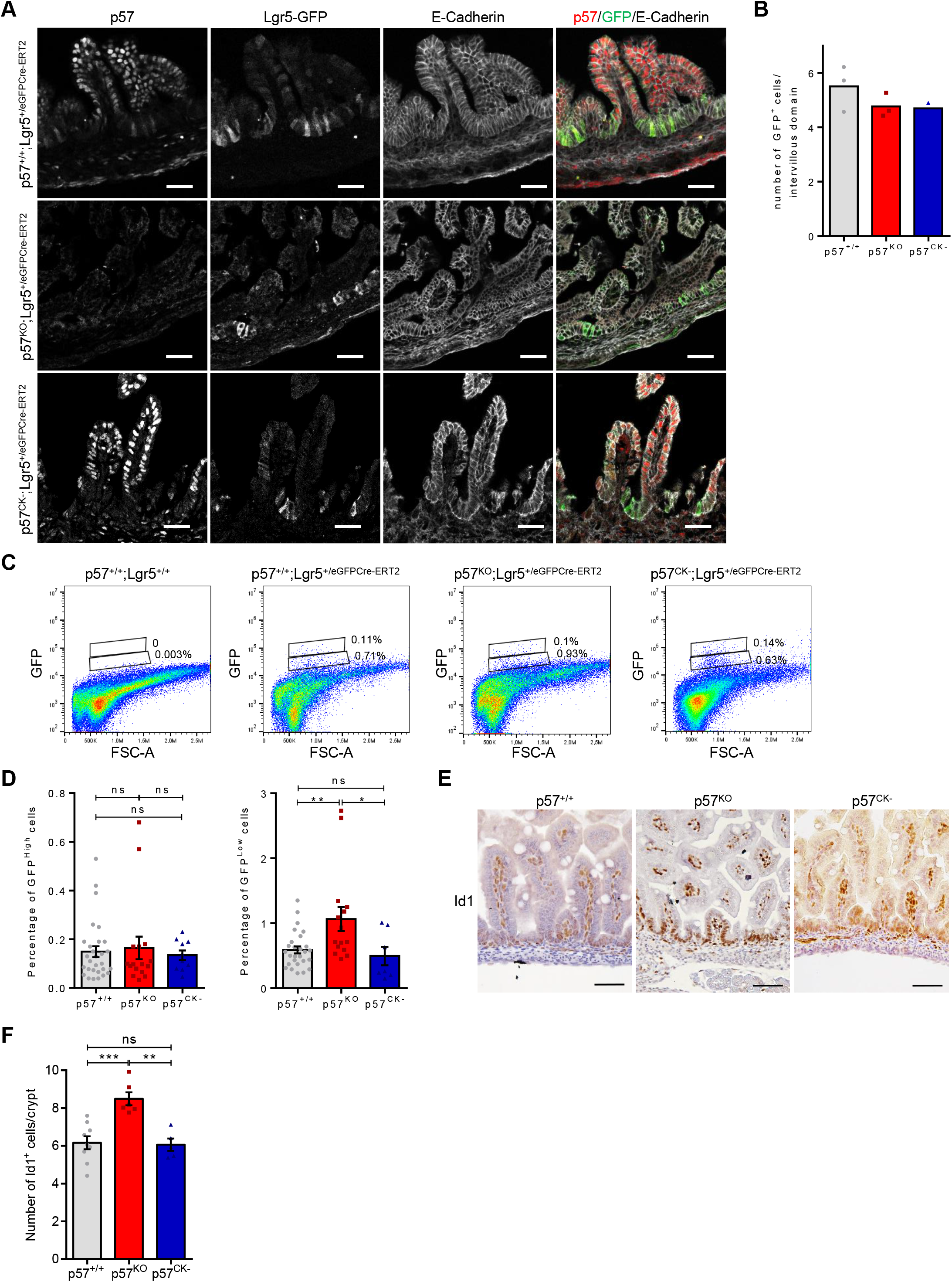
Loss of p57 does not affect Lgr5^+^ ISCs. (A) Representative images of p57^+/+^;Lgr5^+/eGFP-CreERT^, p57^KO^;Lgr5^+/eGFP-CreERT^ and p57^CK-^ ;Lgr5^+/eGFP-CreERT^ intestine sections at E18.5 immunostained for p57 (red), GFP (green) and E-Cadherin (white). Scale bars, 50 µm. (B) Number of Lgr5^+^ (GFP^+^) cells by intervillous domain in p57^+/+^;Lgr5^+/eGFP-CreERT^ (n=5), p57^KO^;Lgr5^+/eGFP-CreERT^ (n=4) and p57^CK-^;Lgr5^+/eGFP-CreERT^ (n=2) intestines counted from immunostained sections as described in A. Graph shows means. (C) Representative dot plots of dissociated intestines from p57^+/+^;Lgr5^+/+^, p57^+/+^;Lgr5^+/eGFP-CreERT^, p57^KO^;Lgr5^+/eGFP-CreERT^ or p57^CK-^;Lgr5^+/eGFP-CreERT^ E18.5 embryos analyzed by flow cytometry for GFP expression. Respective percentages of GFP^High^ (CBCs, top) and GFP^Low^ (TAs, bottom) are indicated for the individuals displayed. (D) Mean percentage of GFP^High^ (CBCs, left panel) and GFP^Low^ (TA cells, right panel) cells from flow cytometry analyses of p57^+/+^;Lgr5^+/eGFP-CreERT^ (n=29), p57^KO^;Lgr5^+/eGFP-CreERT^ (n=16) or p57^CK-^;Lgr5^+/eGFP-CreERT^ (n=10) E18.5 intestines as described in C. Graphs shows means ± SEM. ns: p > 0.05; *: p ≤ 0.05; **: p ≤ 0.01. (E) Representative images of Id1 immunohistochemistry (brown) of intestine sections from p57^+/+^, p57^KO^ and p57^CK-^ E18.5 embryos. Scale bars, 50 µm. (F) Number of Id1^+^ cells per intervillous domain in p57^+/+^ (n=9), p57^KO^ (n=6) and p57^CK-^ (n=5) intestines as described in E. Graph shows means ± SEM. ns: p > 0.05; **: p ≤ 0.01; ***: p ≤ 0.001. See also Figure S2.

### p57 deficiency leads to hyperproliferation of Hopx^+^ reserve ISCs

To label reserve ISCs, p57^KO^ and p57^CK-^ mutant mice were crossed with Hopx^3xFlag-eGFP^ reporter mice in which Hopx is expressed as a Flag-tagged eGFP fusion protein [29, 45] (referred to as p57^+/+^;Hopx^+/3xFlag-eGFP^, p57^KO^;Hopx^+/3xFlag-eGFP^ and p57^CK-^;Hopx^+/3xFlag-eGFP^). Flow cytometry analyses in E18.5 intestines revealed a marked increase of the number of Hopx-GFP^+^ expressing cells in absence of p57 (0.94% ± 0.09) compared to p57^+/+^ (0.56% ± 0.03) and p57^CK-^ (0.39% ± 0.02) mice (Figure 3A, B). The molecular identity of Hopx-GFP^+^ expressing cells was verified by RT-qPCR after sorting and as expected, these cells had increased expression of Hopx, Sox9 and Mex3a [11, 36] (Figure S3A), suggesting that Hopx^+^ cells also act as reserve ISCs in the developing intestine. A modest but consistent enrichment of the CBC markers Lgr5 and Ascl2 was also observed in these cells (Figure S3B), in agreement with previous reports [11, 36, 37]. Finally, Hopx-GFP^+^ cells expressed significant amounts of Atoh1, consistent with the idea that reserve ISCs are in fact progenitors committed to the secretory lineage (Figure S3C) [36]. Reserve ISCs are mostly quiescent, with approximately 20% of cycling cells [25, 27, 29, 46]. To determine if the increased number of Hopx-GFP^+^ cells in p57^KO^ intestine is caused by an exit from quiescence, their proliferation was evaluated by GFP/PCNA co-immunostaining (Figure 3C-E). Consistent with the flow cytometry results, quantification of GFP^+^ cells confirmed the increased number of Hopx-GFP^+^ cells per intervillous domain in p57^KO^ mice (Figure 3C). Moreover, there was a strong increase in the number of PCNA positive Hopx-GFP^+^ cells in p57^KO^ (∼65%), while it was approximately 25% in p57^+/+^ and p57^CK-^ intestines (Figure 3D, E).

**Figure 3:**
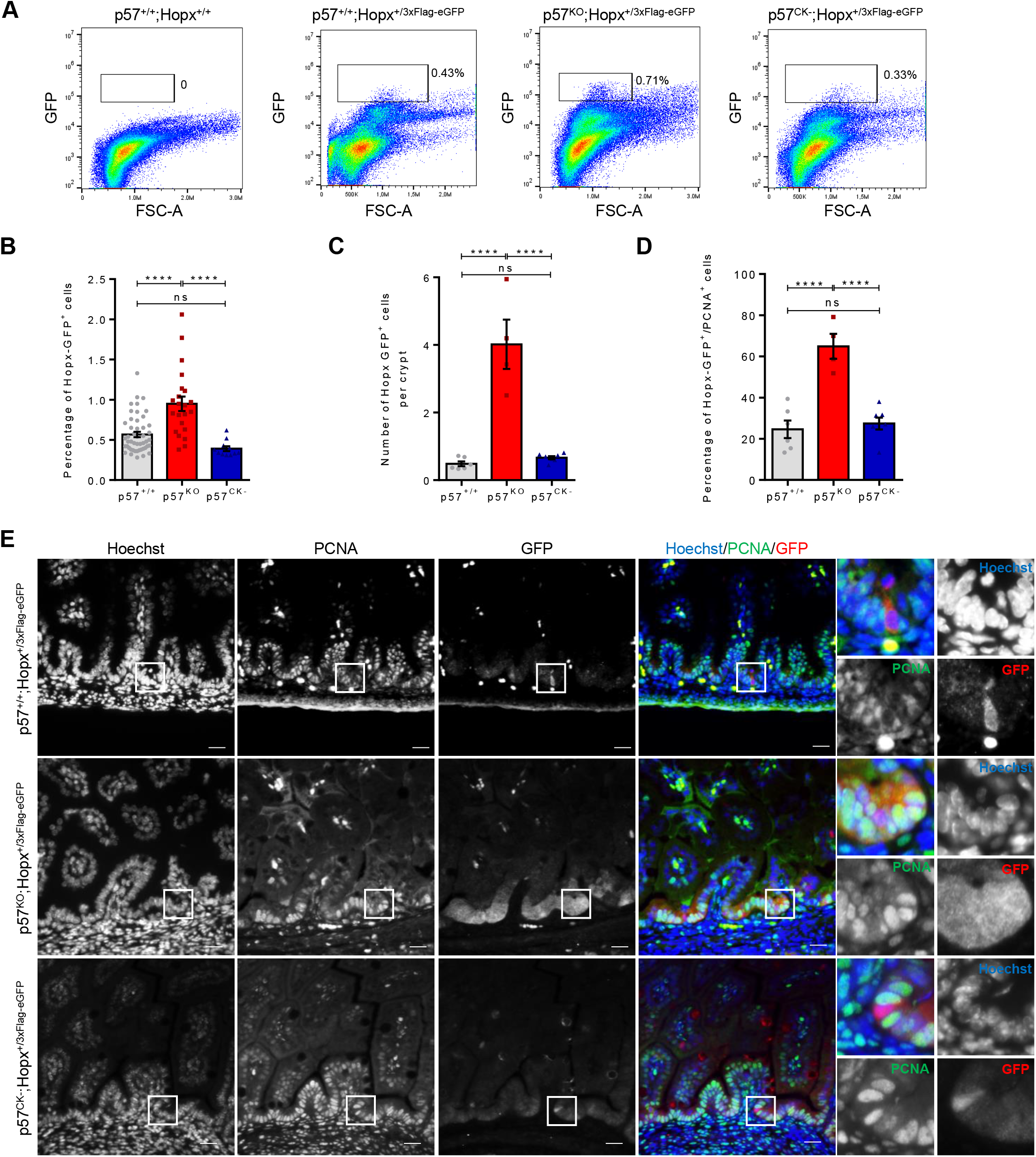
Amplification and exit from quiescence of Hopx^+^ ISCs in p57^KO^ mice. (A) Representative dot plots of dissociated intestines from p57^+/+^;Hopx^+/+^, p57^+/+^;Hopx^+/3xFlag-eGFP^, p57^KO^;Hopx^+/3xFlag-eGFP^ and p57^CK-^;Hopx^+/3xFlag-eGFP^ E18.5 embryos analyzed by flow cytometry for GFP expression. Respective percentage of GFP^+^ cells are indicated for the individuals displayed. (B) Mean percentage of Hopx-GFP^+^ cells in flow cytometry analyses of p57^+/+^;Hopx^+/3xFlag-eGFP^ (n=51), p57^KO^;Hopx^+/3xFlag-eGFP^ (n=23) and p57^CK-^;Hopx^+/3xFlag-eGFP^ (n=14) E18.5 intestines as described in A. Graph shows means ± SEM. ns: p > 0.05; ****: p ≤ 0.0001. (C) Mean number of Hopx-GFP^+^ cells by intervillous domain in p57^+/+^;Hopx^+/3xFlag-eGFP^ (n=6), p57^KO^;Hopx^+/3xFlag-eGFP^ (n=4) and p57^CK-^;Hopx^+/3xFlag-eGFP^ (n=7) E18.5 intestines immunostained for GFP and PCNA as shown in E. Graph shows means ± SEM. ns: p > 0.05; ****: p ≤ 0.0001. (D) Mean percentage of proliferative Hopx-GFP^+^ ISCs, positive for both GFP and PCNA, in intestines as described in C. Graph shows means ± SEM. ns: p > 0.05; ****: p ≤ 0.0001. (E) Representative images of GFP and PCNA immunostaining of p57^+/+^;Hopx^+/3xFlag-eGFP^, p57^KO^;Hopx^+/3xFlag-eGFP^ and p57^CK-^;Hopx^+/3xFlag-eGFP^ E18.5 intestines. DNA was stained with Hoechst 33342. Scale bars, 20 µm. See also Figure S3 and S4.

Sox9 was previously described as an ISC marker, with Lgr5^+^ ISCs being Sox9^Low^ and reserve ISCs Sox9^High^ [30, 31, 47]. To confirm our results, Sox9 immunostaining was used to monitor both ISC populations in function of p57 status (Figure S4). In agreement with the results obtained by genetic labeling of Hopx^+^ and Lgr5^+^ cells, the absence of p57 clearly increased the number of Sox9^High^ cells, but did not affect the number of Sox9^Low^ cells (Figure S4A-C). In contrast, p57^CK-^ mice had the same phenotype as p57^+/+^ mice. Moreover, co-immunostaining for Sox9 and PCNA confirmed that loss of p57 induces proliferation of reserve ISCs, with ∼65% of Sox9^High^ cells expressing PCNA, while only ∼20% of Sox9^High^ cells were PCNA positive in p57^+/+^ and p57^CK-^ intestines (Figure S4D, E), as expected [31, 46]. Overall, these results suggest that p57 plays a critical role in maintaining quiescence in reserve ISCs, independently of CDK inhibition.

### Loss of p57 causes major gene expression changes in Hopx^+^ ISCs

The expression of p57 was evaluated by RT-qPCR in both ISC populations. Consistent with p57 immunostaining (Figure 1G and S1B) showing no signal in lower crypts, p57 was expressed at low levels in Lgr5^+^ ISCs (GFP^High^) and TA cells (GFP^Low^) but was abundant in differentiated (GFP^Neg^) cells (Figure 4A). In contrast, p57 mRNA expression was high in Hopx-GFP^+^ cells (Figure 4B), in agreement with the reported expression of p57 in quiescent Mex3a^High^ or Bmi1^High^ ISCs [11, 12]. To confirm the expression of p57 in +4 reserve ISCs, p57 expression was investigated by immunostaining in E18.5 intestines from p57^+/+^;Hopx^+/3xFlag-eGFP^. In keeping with RT-qPCR results, p57 expression was detected in Hopx-GFP^+^ cells localized around the +4 position (Figure 4C, D). Interestingly, cytoplasmic localization of p57 was observed in some, but not all, Hopx-GFP^+^ cells (Figure 4C), suggesting an active regulation of p57 shuttling in these cells.

**Figure 4:**
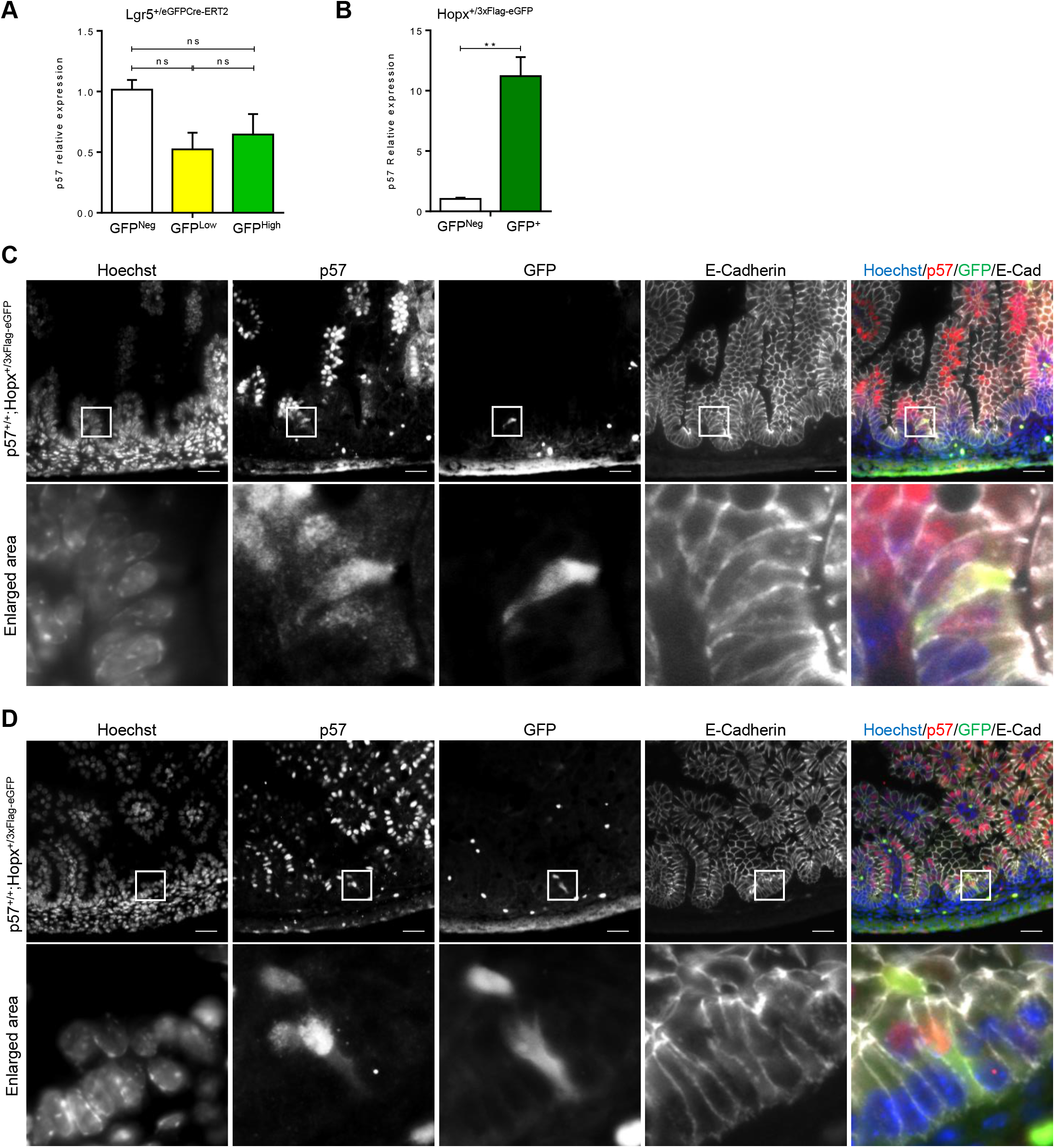
p57 is expressed in Hopx^+^ stem cells. (A, B) Relative p57 expression levels, evaluated by RT-qPCR, in cells sorted based on GFP expression from dissociated crypts from Lgr5^+/eGFP-CreERT^ (A) and Hopx^3xFlag-eGFP^ (B) intestines. p57 mRNA levels were normalized to GAPDH expression and expressed relative to expression in GFP^Neg^ cells. (C, D) Representative images of GFP, p57 and E-Cadherin immunostaining of p57^+/+^;Hopx^+/3xFlag-eGFP^ E18.5 intestines. DNA was stained with Hoechst 33342. Scale bars, 25 µm.

To further define the role of p57 in Hopx^+^ ISCs, RNA-Seq analyses were performed on Hopx-GFP^+^ cells sorted from p57^+/+^ or p57^KO^ E18.5 intestines to compare their bulk RNA profiles. In the entire genome, 21110 genes were expressed at appreciable level (>50 RPKM in total on the 10 samples). Differential DESeq2 analysis revealed that 6111 genes were deregulated in p57^KO^ mice (p<0.05 and absolute Log2 fold change>1), with 3129 genes upregulated and 2982 downregulated (Figure 5A, B and Table S2). We next investigated how p57 status affected the gene expression signature of different intestinal cell populations previously published [11, 48]. Strikingly, loss of p57 resulted in increased expression of genes found in the signature of CBCs and TA cells (Figure 5C, D), supporting the idea that p57^KO^ Hopx^+^ cells exit quiescence and behave like proliferative ISCs. For instance, in Hopx^+^ ISCs, expression of Lgr5, Ascl2, Olfm4, and Id1 approximately doubled in p57^KO^ compared to p57^+/+^ (mean FPKM 5.07 vs 2.62; 3.87 vs 1.95; 102.6 vs 57.42; and 61.53 vs 26.2, respectively). The signature genes of LRCs and Mex3a^High^ cells were also expressed at higher levels in p57^KO^ Hopx^+^ ISCs (Figure 5E, F). Interestingly, p57^KO^ Hopx^+^ cells also expressed higher levels of genes present in differentiated intestinal epithelial cells of both secretory (Enteroendocrine and Goblet) and absorptive (Enterocytes) lineages (Figure S5). This likely reflects the increased number of committed progenitors (TA cells) in absence of p57 and indicates that Hopx^+^ cells are capable of generating both secretory and absorptive progenies. Taken together, these results indicate that in absence of p57, quiescent Hopx expressing cells become proliferative and acquire markers of proliferative Lgr5^+^ ISC population, as well as of committed progenitor cells.

**Figure 5:**
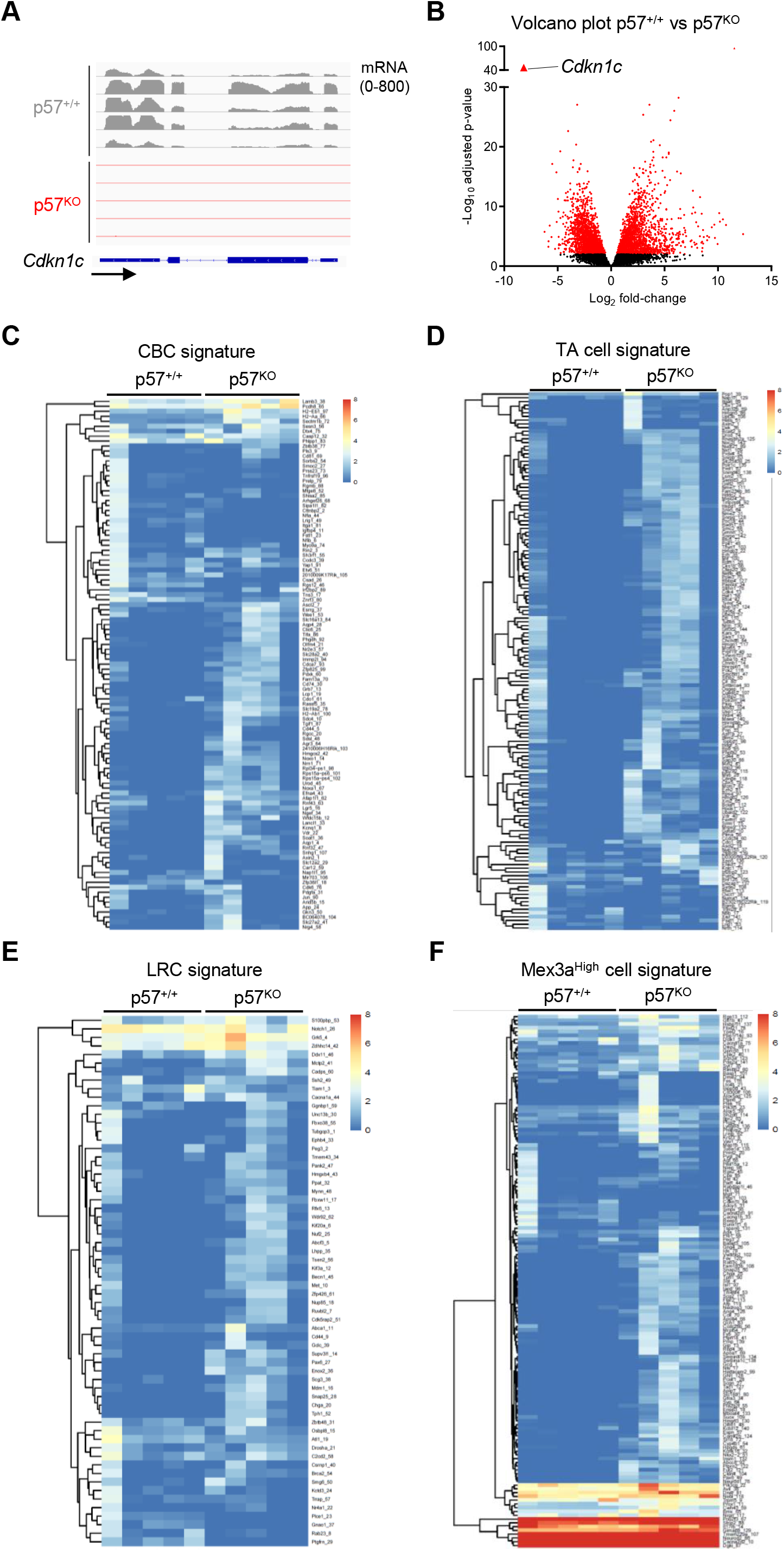
Transcriptomic analyses show major gene expression changes in Hopx^+^ ISCs of p57^KO^ mice. Hopx-GFP^+^ cells were sorted from dissociated crypts from E18.5 p57^+/+^ (n=5) and p57^KO^ (n=5) embryos and submitted to RNA-Seq analysis. (A) Genome Browser image capture for RNA-Seq coverage of p57 mRNA (|Log_2_FC|=-8.7; p adj.=3.9e^-47^). (B) Volcano plot for mRNA expression changes in p57^+/+^ vs p57^KO^ Hopx-GFP^+^ cells as described above. Differential gene expression in function of p57 status was determined by DEseq2 analysis of RNA-Seq. Red dots represent significantly deregulated genes (|Log_2_FC| >1 and adjusted p-value <0.05). (C-F): Heatmaps of Hopx-GFP^+^ ISC RNA-Seq FPKM from p57^+/+^ (n=5) and p57^KO^ (n=5) mice for previously published gene expression signatures of specific intestinal cell populations: CBCs (C), Transit amplifying (TA) cells (D), Label Retaining Cells (LRCs) (E) and Mex3a^High^ cells (F). See also Figure S5.

### p57 interacts with Ascl2 and inhibits its transcriptional activity

In neural progenitors, p57 was reported to promote their maintenance and prevent differentiation by binding to and inhibiting the transcription factor Ascl1 [49]. Ascl2, a paralog of Ascl1, is also a basic helix-loop-helix transcription factor that activates transcription by binding DNA on E-box motifs (CANNTG) [41]. Ascl2 plays a key role in Lgr5^+^ ISC specification and maintenance and its re-expression and activation is required for Lgr5^+^ ISC regeneration following crypt damage [40, 42]. Ascl2 acts synergistically with Wnt signaling to control the expression of key ISC genes, including Lgr5, Sox9 and EphB3 [41]. Here, we tested whether p57 inhibits Ascl2 activity to regulate ISC fate and proliferation.

An interaction between p57 and Ascl2 was evidenced by co-immunoprecipitation (co-IP) in HEK293 cells (Figure 6A, B). This interaction was also detected between p57^CK-^ and Ascl2, indicating that the interaction does not require cyclin-CDK complexes, however, in these experiments, p57^CK-^ binding to Ascl2 was consistently lower than with wild-type p57, suggesting that the point mutations in the cyclin and/or CDK binding domains of p57^CK-^ affect this interaction (Figure 6A, B). Indeed, the interaction of p57 with other transcription factors such as MyoD or b-Myb was previously reported to take place within the cyclin-CDK binding region of p57 [50-52]. The p57/Ascl2 interaction was confirmed by proximity ligation assays (PLA) on endogenous proteins in SW480 cells (Figure 6C). The effect of p57 on Ascl2 transcriptional activity was determined in transcription reporter assays in which a destabilized GFP (half-life ∼1 h) under the control of the Sox9 or Lgr5 promoters (Figure 6D) was co-transfected in HEK293 cells with Ascl2 and increasing amounts of p57 or p57^CK-^. Results were normalized to the activity of the GAPDH promoter obtained in the same conditions. Transfection of increasing amounts of p57 decreased Sox9 and Lgr5 promoter activity, indicating a significant inhibition of Ascl2 transcriptional activity, while no statistically significant effect was observed when the Phox2A promoter, an Ascl1 target [49], was used (Figure 6E). Similar results were obtained when p57^CK-^ was expressed (Figure 6F), indicating that repression of Ascl2 activity is a CDK-independent process. Finally, monitoring of the expression of 21 known Ascl2 target genes [42] in the RNA-Seq from sorted Hopx-GFP^+^ cells indicated that 11 of them were significantly upregulated in p57^KO^ compared to p57^+/+^ Hopx^+^ cells (Figure 6G), comforting the idea that p57 acts as a transcriptional repressor towards Ascl2. Thus, these data indicate that p57 can bind to and inhibit Ascl2 transcriptional activity in a CDK-independent manner.

**Figure 6:**
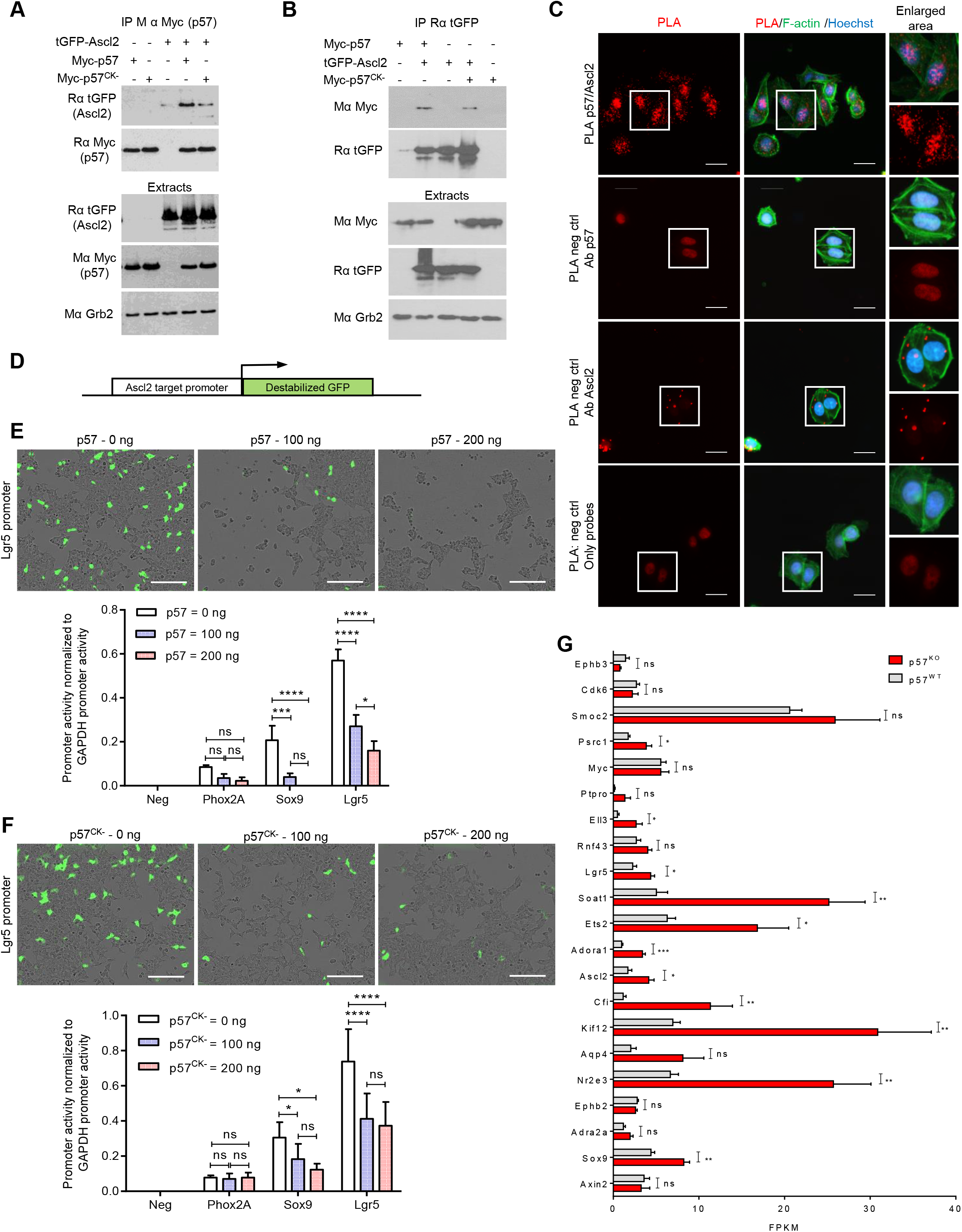
p57 binds to and inhibits Ascl2. (A) p57 or p57^CK-^ was immunoprecipitated with Myc antibodies from HEK293 cells transfected with tGFP-Ascl2 and/or Myc-p57 or Myc-p57^CK-^ and the presence of Ascl2 was assessed by immunoblotting (n=3). Amounts of transfected proteins in lysates are shown. Grb2 was used as loading control. (B) Ascl2 was immunoprecipitated with tGFP antibodies from HEK293 cells transfected with tGFP-Ascl2 and/or Myc-p57 or Myc-p57^CK-^ and the presence of p57 was determined by immunoblot (n=3). Amounts of transfected proteins in lysates are shown. Grb2 was used as loading control. (C) Proximity Ligation Assay (PLA) using p57 and Ascl2 antibodies on endogenous proteins in SW480 cells. F-actin was stained with phalloidin and DNA with Hoechst 33342. Scale bars, 50 µm. Representative images from three independent experiments. (D) Schematic of a pZsGreen reporter construct in which a destabilized GFP (½ life ∼ 1 h) is cloned downstream of a promoter to measure promoter activity. (E-F) HEK293 cells were co-transfected with 100 ng of a reporter construct (pZsGreen DA [no promoter, negative control], or pZsGreen pGAPDH [positive control], or pZsGreen pPhox2a [Ascl1 target], or pZsGreen pSox9, or pZsGreen pLgr5 [Ascl2 targets]), 100 ng of an Ascl2 expression vector, and the indicated amounts of a p57 (E) or p57^CK-^ (F) expression vector. The number of fluorescent cells was quantified using an Incucyte FLR. Scale bars, 200 µm. Graphs represent the mean fluorescent object density normalized to that of GAPDH in the same condition from four independent experiments. Graphs show means ± SEM. ns: p > 0.05; *: p ≤ 0.05; ***: p ≤ 0.001; ****: p ≤ 0.0001. (G) Expression levels FPKM of Ascl2 target genes from [42] in RNA-Seq data from p57^+/+^ and p57^KO^ Hopx-GFP^+^ ISCs from Figure 4. Graphs show means ± SEM. ns: p > 0.05; *: p ≤ 0.05; ***: p ≤ 0.001.

### p57 participates in the recruitment of a corepressor complex to Ascl2 target gene promoters

We previously identified HDAC7 as a potential p57 binding partner in the p57 interactome [5]. HDAC7 is a class II histone deacetylase (HDAC) that represses transcription via the formation of a repressor complex, notably with the scaffold protein mSin3a [53]. Interactions between p57 or p57^CK-^ and HDAC7 (Figure 7A) or mSin3a (Figure 7B) were detected by co-IP in HEK293 cells. Again, p57^CK-^ appeared to interact more weakly with these partners than wild-type p57, especially with mSin3A, indicating that (i) p57 binding to these partners is independent of cyclin/CDK complexes and (ii) that the N-terminal cyclin/CDK binding region may be involved in these interactions. The formation of a protein complex between p57, HDAC7 and mSin3a was validated by co-IP on endogenous proteins in SW480 cells (Figure 7C, D), and these interactions were confirmed by PLA experiments as well (Figure 7E). Similarly, co-IP experiments revealed that Ascl2 also binds to HDAC7 and mSin3a in HEK293 cells overexpressing these proteins (Figure S6A), and these interactions were confirmed by PLA on endogenous proteins in SW480 cells (Figure S6B).

**Figure 7:**
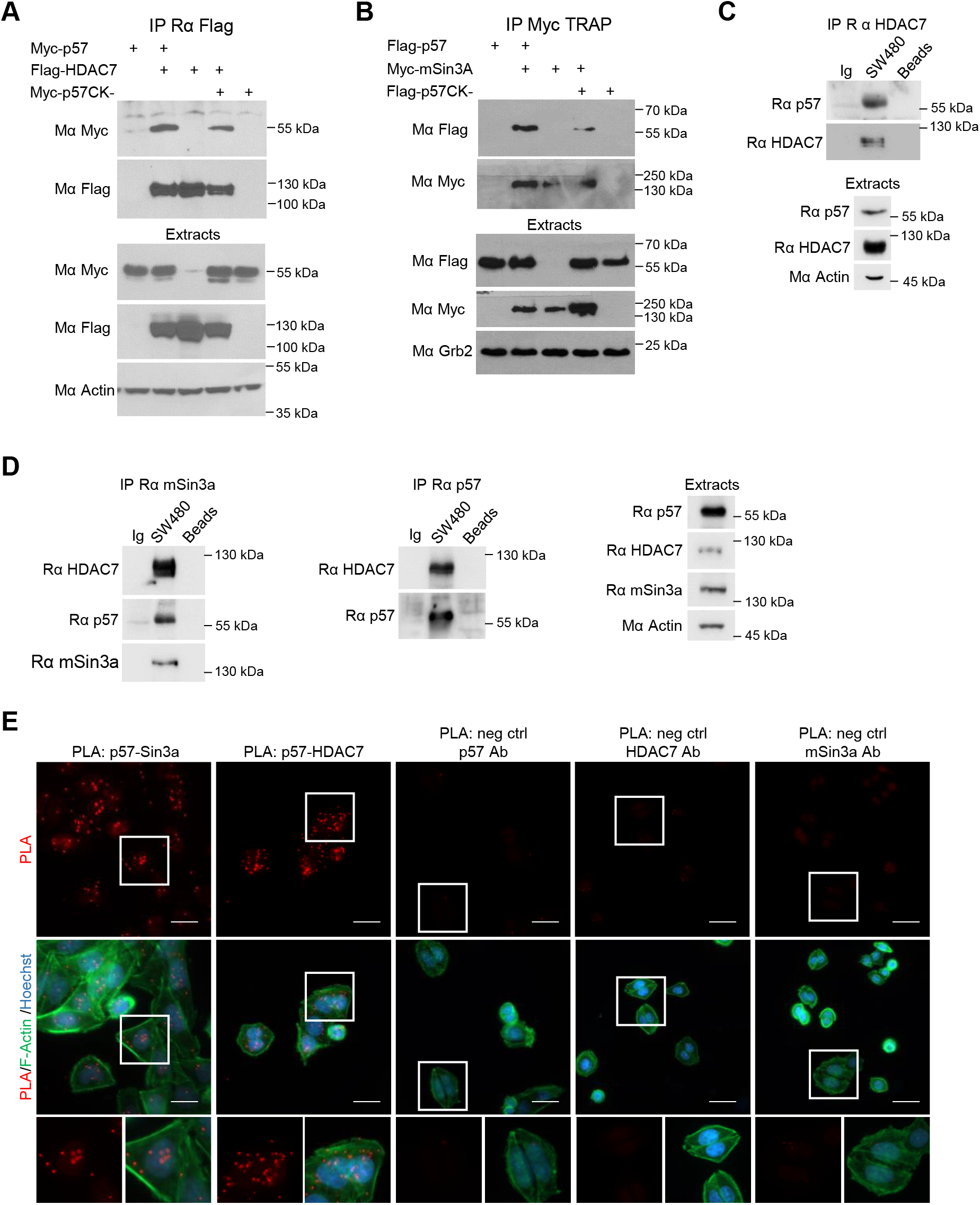
p57 forms a co repressor complex with HDAC7 and mSin3a. (A, B) HDAC7 (A) or mSin3a (B) were immunoprecipitated from HEK293 cells transfected with Flag-HDAC7 or Myc-mSin3a and/or Flag-p57 and/or Flag-p57^CK-^ and the presence of p57 was assessed by immunoblotting (n=3). Amounts of transfected proteins in lysates are shown. Actin (A) or Grb2 (B) were used as loading control. (C-D) Endogenous HDAC7 (C), mSin3a and p57 (D) were immunoprecipitated from SW480 cells lysates and the presence of endogenous p57, HDAC7 and/or mSin3a was assessed by immunoblotting. Protein A beads alone and control IgG were used as negative control. Respective levels of each protein in lysates are shown. β-actin was used as loading control (n=3). (E) PLA for p57/mSin3a proximity and p57/HDAC7 proximity in SW480 cells using p57, mSin3a and/or HDAC7 antibodies. F-actin was stained with phalloidin and DNA with Hoechst 33342. Scale bars, 50 µm. Representative images of three independent experiments. See also Figure S6

Finally, the presence of p57 and the corepressor complex subunits on Ascl2 target gene promoters was investigated by chromatin immunoprecipitation (ChIP) in SW480 cells. As expected, Ascl2 was present on the promoters of Lgr5, Sox9 and EphB3 (Figure 8A and S7A) [42], and p57, HDAC7 and mSin3a were also detected on the same sequences of these Ascl2 target promoters (Figure 8B-D and S8B-D). To confirm that p57 is recruited to DNA by Ascl2, pull-down assays using streptavidin beads bound to biotinylated oligonucleotides containing the E-box of the Lgr5 promoter or a control oligonucleotide with point mutations in the E-Box were performed on HEK293 cells transfected with Ascl2 and/or p57 (Figure 8E) or p57^CK-^ (Figure 8F). p57 and p57^CK-^ required DNA-bound Ascl2 and a functional E-Box to bind the beads (Figure 8E, F). Similar experiments with cells also expressing HDAC7 or mSin3A indicated that these corepressor proteins bound to the E-box-containing oligonucleotide but not to the E-Box mutant oligonucleotide (Figure 8G, H).

**Figure 8:**
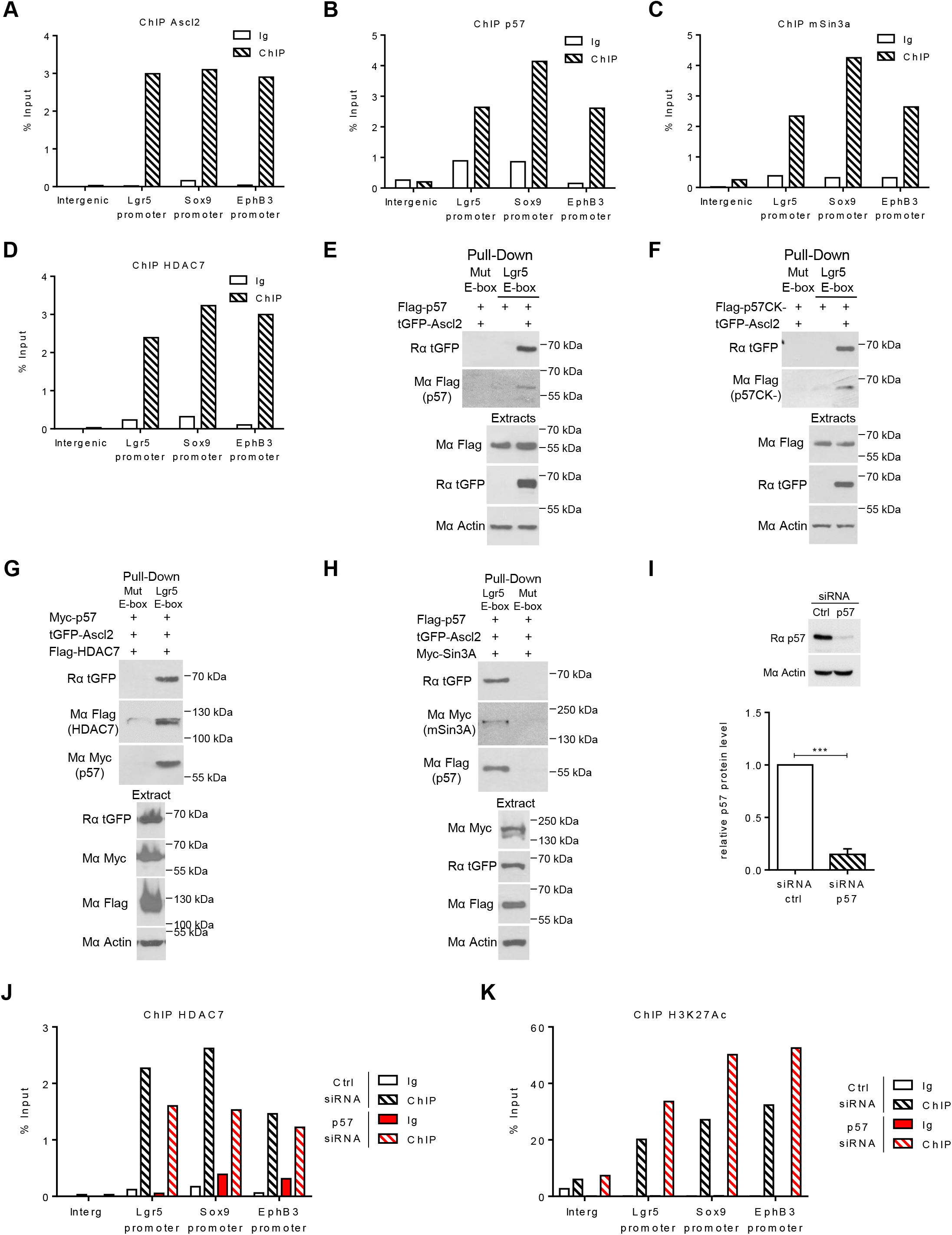
p57 participate in the recruitment of a co-repressor complex on Ascl2 target promoters. (A-D) ChIP experiments to determine the presence of a co-repressor complex on Ascl2 target promoters. ChIP using antibodies against Ascl2 (A), p57 (B), mSin3a (C) and HDAC7 (D), or isotype control, were followed by qPCR with primers specific for the Lgr5, Sox9 or EphB3 promoters or for an unrelated intergenic region as negative control. Results are expressed as a percentage of input. Graphs display the results of one representative assay from three independent experiments and the two other experiments are provided in Figure S7A-D. (E-H) Pulldown assays with streptavidin beads bound to biotinylated oligonucleotides derived from the E-box containing region of the Lgr5 promoter or to oligonucleotides in which point mutations disrupt the E-box as negative control were performed on cell lysate from HEK293 transfected with Ascl2 and p57 (E) or p57^CK-^ (F), p57, Ascl2 and HDAC7 (G) or mSin3a (H). The presence of the different proteins was assessed by immunoblotting. Corresponding levels of transfected proteins in the extracts are shown. β-actin was used as loading control. (I) p57 silencing in SW480 cells. SW480 cells were transfected with control or p57 siRNA for 48 h and subjected to immunoblot for p57. β-actin was used as loading control. Graph shows mean p57 silencing ± SEM from four independent experiments. ***: p ≤ 0.001. (J) ChIP with HDAC7 antibodies or control IgG in SW480 cells transfected for 48 h with control or p57 siRNA followed by qPCR with primers specific for the Lgr5, Sox9 and EphB3 promoters or for an unrelated intergenic region as negative control. Results are expressed as a percentage of input. Graph displays the results of one representative assay from three independent experiments and the two other experiments are provided in Figure S7E. (K) ChIP with H3K27-Ac antibodies or control IgG in SW480 cells transfected for 48 h with control or p57 siRNA followed by qPCR with primers specific for the Lgr5, Sox9 and EphB3 promoters or for an unrelated intergenic region as negative control. Results are expressed as a percentage of input. Graph displays the results of one representative assay from four independent experiments and the two other experiments are provided in Figure S7F. See also Figure S7.

To determine whether p57 mediates the recruitment of the corepressor complex proteins to Ascl2 target genes, ChIPs were performed after p57 silencing with siRNAs (Figure 8I). The knockdown of p57 markedly decreased the amount of HDAC7 present on Ascl2 target gene promoters in SW480 cells (Figure 8J and S8E). Consistent with the idea that p57 participates in the recruitment of a corepressor complex with histone deacetylase activity to Ascl2 target gene promoters, increased histone H3 lysine acetylation (H3K27-Ac) was observed at these promoters when p57 expression was silenced, suggesting a derepression of chromatin (Figure 8K and S8F). Overall, these results suggest that p57 is able to inhibit Ascl2 transcriptional activity by participating in the recruitment of a corepressor complex containing HDAC7 and mSin3a to Ascl2 target gene promoters, thereby regulating ISC fate and proliferation.

## Discussion

The entire intestinal epithelium is renewed every few days by Lgr5^+^ ISCs residing at the bottom of the crypts [15]. In case of genotoxic or cytotoxic damage, Lgr5^+^ ISCs are lost and must be rapidly regenerated to maintain epithelial integrity [14]. Whether Lgr5^+^ ISCs are regenerated by dedifferentiation of committed progenitors that re-enter the niche or by dedicated reserve stem cells that are normally quiescent is still debated [22, 23, 25, 29, 32, 36, 40, 47]. We found that during late development, loss of p57 expression triggers an expansion of the dividing compartment in intestinal crypts, with an amplification of TA cells and of a Hopx^+^ cell population previously described as reserve ISCs in adult mice [45], while the number of Lgr5^+^ ISCs was not affected. In absence of p57, Hopx^+^ ISCs become proliferative instead of being mostly quiescent and cell cycle re-entry is associated with a derepression of Ascl2 activity and increased expression of its target genes. This is consistent with a recent report showing that Ascl2 re-expression in committed progenitors is required for Lgr5^+^ ISC regeneration after damage [40]. Thus, during intestinal development, p57 appears to limit the stem cell niche and suppress the proliferative ISC phenotype by repressing Ascl2 activity in Hopx^+^ cells that are located just above the niche. Similarly, a recent study reported that in adult mice, p57 is specifically expressed in Bmi1^+^ reserve ISCs and is involved in maintaining their quiescence, although the mechanism involved was not investigated [12].

RNA-Seq analyses of sorted Hopx^+^ cells from E18.5 intestines show that p57 loss shifts their gene expression signature towards that of CBCs and a number of cell cycle regulatory genes, including Cyclins A2, B1 and B2, CDK4, Cdc25a and Cdc20, are upregulated, consistent with re-entry in the cell cycle. In addition, differentiation markers for all intestinal cell types are also present in this sorted cell population, suggesting that these Hopx^+^ cells are indeed *bona fid*e ISCs capable of giving rise to progenitors committed towards all intestinal lineages. Importantly, intestine development and ISC populations were normal in p57^CK-^ mice, indicating that p57-mediated cyclin-CDK inhibition is not essential to maintain intestinal epithelium homeostasis and quiescence outside of the crypt. Quiescence of differentiated cells is probably enforced by other CKIs such as p27, which has a similar expression pattern as p57 in the intestine [10]. In Hopx^+^ expressing cells, our data suggest that p57 maintains quiescence by inhibiting the transcriptional activity of Ascl2, thereby repressing the Lgr5^+^ ISC phenotype.

p57^KO^ mice display numerous development anomalies, however the penetrance and presentation of these phenotypes is very variable and affected by the genetic background [3, 5, 9]. For instance, only 30-40% of p57^KO^ animals display intestinal truncation or shortening [3, 5, 9]. Nevertheless, all p57^KO^ animals we examined exhibited histological and molecular alterations, with hyperproliferation in the crypts and gene expression profiles modifications in Hopx^+^ expressing cells regardless of the presence of intestinal shortening. This absence of correlation between macroscopic phenotypes and cellular/molecular defects could be an illustration of this variability in phenotype presentation and severity. The reasons for this variability remains to be elucidated. Of note, the presentation and severity of phenotypes in BWS patients is also very variable [6, 7].

Several studies have described the ability of p57 to regulate transcription by distinct mechanisms. For instance, p57 binds to and promotes MyoD activity by preventing CDK-mediated phosphorylation and targeting for degradation of MyoD, as well as by another unknown mechanism independent of CDKs [50]. Conversely, p57 was found to bind to and inhibit the transcriptional activity of Ascl1 independently of CDKs, although the underlying mechanism was not investigated [49]. p57 may also control transcription by regulating RNA Polymerase II C-terminal domain (CTD) phosphorylation in an E2F1-dependent manner [54]. More recently, p57 was found to promote c-Jun and FHL2 mediated transcription by competing with HDAC1 and HDAC3 for binding to these transcription factors, thereby relieving their inhibition by HDACs [55, 56]. Interestingly, while the N-terminus of p57 acted as a transactivator, presumably by competing with HDACs, the C-terminus of p57 repressed c-Jun activity, suggesting a complex regulation that may depend on the binding partners of p57 and/or on posttranslational modifications [55]. Herein, we found that p57 could interact with HDAC7 and with the corepressor mSin3a, in a CDK independent manner, suggesting that p57 may function as a transcriptional corepressor by recruiting a repressor complex to specific promoters, in a manner similar to p27 [57]. In agreement with this idea, p57 is present on chromatin at Ascl2 target promoters and appears to control the recruitment of these corepressors to promoters. Indeed, p57 knockdown resulted in decreased presence of HDAC7 on Ascl2 target gene promoters, as well as in increased Histone H3 acetylation levels, which is consistent with derepression of these promoters, a possibility that was confirmed by RNA-Seq analyses.

Interestingly, two transcriptional corepressors, RUNX1T1 (MTG8) and CBFA2T3 (MTG16) were recently found to be expressed in cells at the +4/+5 positions and to control the fate of these progenitors/stem cells, directing them towards differentiation in the enterocyte lineage [58]. *Runx1t1* knockout and *Runx1t1*/*Cbfa2t3* double knockout mice die shortly after birth and exhibit intestinal shortening [58]. This is accompanied by increased proliferation in the crypts and increased expression of Lgr5^+^ ISC signature genes [58]. RUNX1T1 and CBFA3T2 act as transcriptional corepressors by binding to mSin3a, several HDACs and the N-CoR and SMRT complexes [59]. Given the phenotypic similarities of the *Runx1t1* and *Runx1t1*/*Cbfa2t3* knockout mice with those of p57^KO^ mice, it would be interesting to test if RUNX1T1 and/or CBFA2T3 are present in the corepressor complex in which p57 is involved. The chromatin state of Lgr5^+^ ISCs and committed progenitors are very similar, although the promoters/enhancers of lineage restricted genes are selectively open in the latter, which may be reversed during their dedifferentiation into Lgr5^+^ ISCs [60-62]. This suggests that the cellular plasticity in the intestinal epithelium and interconversion of ISCs does not require broad modifications of the chromatin state, but may be mediated by modulating the activity of specific transcription factors, such as Ascl2, and/or by epigenetic modifications of a limited number of cis-acting elements [40, 61]. Our data suggest that p57 acts as a key regulator of Ascl2 activity and participates in modulating the permissive state of chromatin at Ascl2-responsive promoters/enhancers. Thus, controlling p57 levels along the crypt-villus axis may constitute a powerful means to regulate cell fate in the intestinal epithelium.

Ascl2 is a target of the Wnt/LEF1/β-catenin pathway whose expression is frequently upregulated in colorectal cancers [42, 63, 64]. Ascl2 was also involved in treatment resistance in colorectal cancer cells [65]. Conversely, *CDKN1C* is a putative tumor suppressor gene and epigenetic silencing of p57 is observed in colorectal cancers, correlating with poor prognosis [66-69]. Moreover, a recent study highlights the role of p57 in maintaining quiescence of colorectal cancer stem cells, conferring resistance to chemotherapeutic agents to these cells [70]. Our results could provide the molecular basis explaining the possible tumor suppressor role of p57 in colorectal cancer. Indeed, p57 silencing may contribute to the derepression of Ascl2 activity in colorectal cancer cells and in the acquisition of cancer stem cell characteristics. Overall, our data suggests that p57 regulates intestinal development by maintaining quiescence of Hopx^+^ reserve ISCs in a CDK-independent manner, by repressing the transcriptional activity of Ascl2 via the recruitment of a corepressor complex. Thus, p57 appears to be a key regulator participating in the control of ISC fate and proliferation.

## Materials and Methods

### Antibodies, Reagents and plasmids

Mouse anti-PCNA (PC10, sc-59), mSin3a (G-11, sc-5299), HDAC7 (A-7, sc-74563), Myc (9E10, sc-40), OctA-probe (Flag, H5, sc-166355), Actin (C4, sc-17778), and rabbit anti-p57 (H-91, sc-8298) and Myc (A-14, sc-789) were purchased from Santa Cruz Biotechnology. Mouse anti-Ascl2 (8F1, MAB4417) and rabbit anti-Sox9 (AB5535) and p57 (06556) were purchased from Merck Millipore. Mouse anti-β Actin (A2228) and rabbit anti-Ascl2 (SAB1305026), HDAC7 (SAB4502143) and p57 (P0357) were purchased from Sigma-Aldrich. Rabbit anti-mSin3a (PA1-870), HDAC7 (PA5-29937) and Ki-67 (SP6, RM-9106-S1) were purchased from Thermo Scientific. Rabbit anti-cleaved caspase 3 (#9664), Olfm4 (#39141), p57 (#2557S), Flag (#3724) and HDAC7 (#10831) were purchased from Cell Signalling Technology. Rabbit anti-turboGFP (AB513) was purchased from Evrogen. Goat anti-GFP (GTX26673) was purchased from Genetex. Rabbit anti-Id1 (BCH-1/#37-2) was purchased from BioCheck. Mouse anti-Grb2 (610112) and E-cadherin (610182) were purchased from BD Transduction Laboratories. Anti-histone H3 acetyl K27 (Ab4729) was purchased from Abcam. Myc-Trap agarose (yta-20) beads were purchased from Chromotek. Secondary antibodies against whole Ig or Ig light-chain conjugated to horseradish peroxydase (715-035-150, 711-035-152); Secondary antibodies against mouse Ig conjugated to Cy2, Cy3 and Cy5 (715-225-150, 715-175-150, 715-165-150, respectively), anti-rabbit Cy2 and Cy3 (715-255-152, 711-165-152, respectively) and anti-goat Cy2 and Cy3 (705-225-147, 705-165-147, respectively) were purchased from Jackson ImmunoResearch. HRP conjugated antibodies were used at 1/10000 and Cyanine-conjugated antibodies were used at 1/400. Phalloidin-Fluoprobes-495 was purchased from Interchim (FP-47548A). Control siRNA (sc-108727) and siRNA anti human p57 (sc-37751) were purchased from Santa Cruz Biotechnologies.

Myc-tagged p57 and p57^CK-^ in pcDNA3.1+ Hygro vector (Invitrogen) were described previously [5]. The coding sequence of murine Ascl2 fused to turbo-GFP (pCMV6-Ascl2-tGFP, MG203459) was purchased from Origene and subcloned in pcDNA3.1+ Hygro. pcDNA3.1+ HDAC7-Flag was a gift from Eric Verdin (Addgene #13824)[71]. pCS2+MT-mSin3a was a gift from Robert Eisenman (Addgene #30452). The transcription reporter vectors pZs-Green-pGAPDH, pZs-Green-pLgr5, pZs-Green-pSox9 and pZs-Green-pPhox2a were generated by cloning the promoter sequences of GAPDH, Sox9, Lgr5 and Phox2a, purchased from GeneCopoeia (Ref. GAPDH-PF02, HPRM25638-PF02, HPRM13362-PF02 and HPRM13365-PF02, respectively) in the pZs-Green1-DR vector (Clontech).

### Mouse models

p57^KO^ and p57^CK-^ mice have been described previously [5, 9]. Mice were maintained in a 129S4/C57BL6J genetic background. They were crossed with Lgr5^+/eGFP-CreERT^ (JAX #008875) or homozygous Hopx^3xFlag-GFP^ (JAX #029271) mice maintained in a C57BL6J genetic background. Mouse husbandry and procedures were performed in accordance with EU and national regulation (Protocol Authorization APAFIS #31318-202104121433391).

The *Cdkn1c* gene is paternally imprinted and p57 is expressed only from the maternal allele. Therefore, maternal inheritance of the mutant allele (p57^+/-m^ or p57^+/CK-m^) causes phenotypes similar to homozygous mutations [4, 5]. To simplify notations, p57^+/-m^ and p57^-/-^ mice are referred to as p57^KO^, and p57^+/CK-m^ and p57^CK-/CK-^ animals are referred to as p57^CK-^.

### Cell culture and transfection

HEK293 cells were grown in DMEM 4.5 g/L glucose (D6429, Sigma) supplemented with 10% Fetal Bovine Serum (FBS) (10270106, Life Technologies), 0.1 mM non-essential amino acid (M7145, Sigma) and 2 µg/mL penicillin/streptomycin (P4333, Sigma). SW480 cells were cultivated in RPMI 2 g/L glucose (R8758, Sigma) supplemented with 10% FBS, 0.1 mM non-essential amino acid, 2 µg/mL penicillin/streptomycin and 2.5 g/L D-Glucose (G7021, Sigma). All cells were grown at 37°C and 5% CO_2_. HEK293 cells were transfected by the calcium phosphate method 24 h prior to lysis. siRNA transfection was performed 48 h before experiments using Interferin (Polyplus transfection) and 50 nM of the indicated siRNA according to the manufacturer’s instructions.

### Histology

Intestines were collected at E18.5 and fixed in 10% neutral buffered formalin for 2 h at 4°C. For cryosectioning, tissues were incubated in 30% sucrose overnight at 4°C and embedded in OCT (LAMB/OCT, Thermo Scientific). Sections of 12-µm thickness were cut on a cryotome (Cryostat Leica CM1950), placed on Superfrost Plus glass slide (Thermo Scientific) and kept at -20°C. Sections were thawed, rehydrated and blocked in PBS-10% Donkey Serum. Tissues were incubated with primary antibodies overnight at 4°C, washed three times 5 min in PBS 0.1% Triton X-100, then incubated with corresponding secondary antibodies conjugated to Cy2 or -3 for 30 min at room temperature. DNA was stained with Hoechst 33342 (0.1 µg/mL) and slides were mounted with Gelvatol (20% v/v glycerol, 10% w/v polyvinyl alcohol, 70 mM Tris, pH 8). Images of cryosections were acquired on an inverted Leica Sp8 confocal microscope or an inverted spinning disk DMI8 Leica microscope equipped with a confocal Yokogawa head CSI-X1-M1N, and processed with the Fiji Software.

Tissues embedded in paraffin were sectioned on a Microm Microtech microtome and serial 5 µm sections were cut and placed on Superfrost Plus slides. Tissues were deparaffinized and stained with hematoxylin and eosin or used for immunostaining. Intestine sections were rehydrated and antigens were unmasked for 30 min in a steamer with sodium citrate (10 mM, pH 6) (Ki67, GFP, p57, PCNA, cleaved caspase 3) or High pH (H-3301, Vector Laboratories) (Olfm4, Sox9, Id1, PCNA) solutions. Slides were washed twice in PBS 0.2% Triton X-100 and once in PBS and then blocked for 0.5 to 2 h in PBS 0.2% Triton X-100, 10% Donkey Serum, 3% BSA solution at room temperature in a humid chamber. Sections were incubated with primary antibodies overnight at 4°C or 1 h at 37°C and washed three times 5 min in PBS 0.2% Triton X-100. For immunofluorescence, sections were incubated with secondary antibodies conjugated to Cy2 and/or Cy 3 for 30 min at 37°C, and washed three times 5 min in PBS. DNA was stained with Hoechst 33342 (0.1 µg/mL) in the first wash and slides were mounted with Gelvatol. For immunohistochemistry, sections were incubated with anti-Rabbit ImmPress Reagent (MP-7401, Vector Laboratories) for 30 min at 37°C, washed three times 5 min in PBS 0.2% Triton X-100 and staining was visualized with the chromogen 3’3’-diaminobenzidine (ImmPACT DAB, SK-4105, Vector Laboratories). Slides were then dehydrated and mounted with Permount mounting medium (SP15-500, Fisher Scientific). Images of paraffin sections were acquired on a Nikon 90i Eclipse microscope using a Digital DS-Fi1 camera (Nikon) or DS-Qi2 HQ Camera (Nikon) and the NIS Element BR Software.

### Flow cytometry

Intestines were dissected at E18.5, minced with a razor blade and incubated with Dispase (1 U/mL) (D4693, Sigma) and DNase I (0.1 mg/mL) (1307, Euromedex) in PBS for 10 min at 37°C under agitation. Single cell suspensions were collected, filtered through a 40-µm mesh and washed once in PBS after centrifugation for 10 min at 2000 rpm. Pelleted cells were resuspended in PBS 2% FBS and dead cells were stained with 7AAD (0.5 µg/mL) (A9400, Sigma) for 30 min at 4°C. Flow cytometry analyses were performed on a Cytoflex S cytometer (Beckman Coulter) and data were analyzed with the FlowJo software. To monitor GFP populations in reporter mice, a gate was drawn on single live cells. Single cells were identified using Forward Side Scatter and live cells using 7AAD staining (7AAD^-^). All analyses were carried out genotype blind.

### Cell sorting

Intestinal crypts were purified as previously described [72]. Briefly, small intestines were dissected and opened longitudinally. The tissue was incubated with HBSS 30 mM EDTA pH 7.4 for 15 min at 37°C. The epithelium was detached from the mucosa by vigorous shaking and the epithelial fraction was incubated in 10 mL HBSS supplemented with DNase I (0.1 mg/mL) and Dispase (1 U/mL) for 20 min at 37°C. Before sorting, a single cell preparation was obtained by centrifugation and resuspension in HBSS 5% FBS and filtering through a 40-µm sieve. Cells were stained with 7AAD (0.5 µg/mL) to exclude dead cells. Single GFP^+^ 7AAD^-^ cells were sorted directly in trizol using a FACSAria Fusion cytometer (Becton Dickinson) for subsequent RNA extraction.

### Reverse transcriptase and qPCR

Cells were lysed in TriReagent (Sigma, T9424) and RNA was extracted according to the manufacturer’s protocol. RNA integrity was verified on 0.8% agarose gel and quantified using a Nanodrop spectrophotometer. cDNA was synthetized using SuperScript IV (Thermo Fisher) reverse transcriptase according to the supplier’s instructions with 1 µg RNA. A negative control without reverse transcriptase was performed for each experiment. qPCR reactions were carried out on a Biorad CFX94 apparatus using Supermix SsoFast EvaGreen (Biorad) with primers at a final concentration of 500 nM and a quantity of cDNA corresponding to 5 ng of RNA per reaction. Ct values were generated with BioRad CFX96 Real-Time PCR software and data were analyzed and normalized using the 2^-ΔΔCt^ method with GAPDH as housekeeping gene. All Ct were expressed relative to expression in the GFP^Neg^ population in the same experiment. The primers used are presented in the following table.

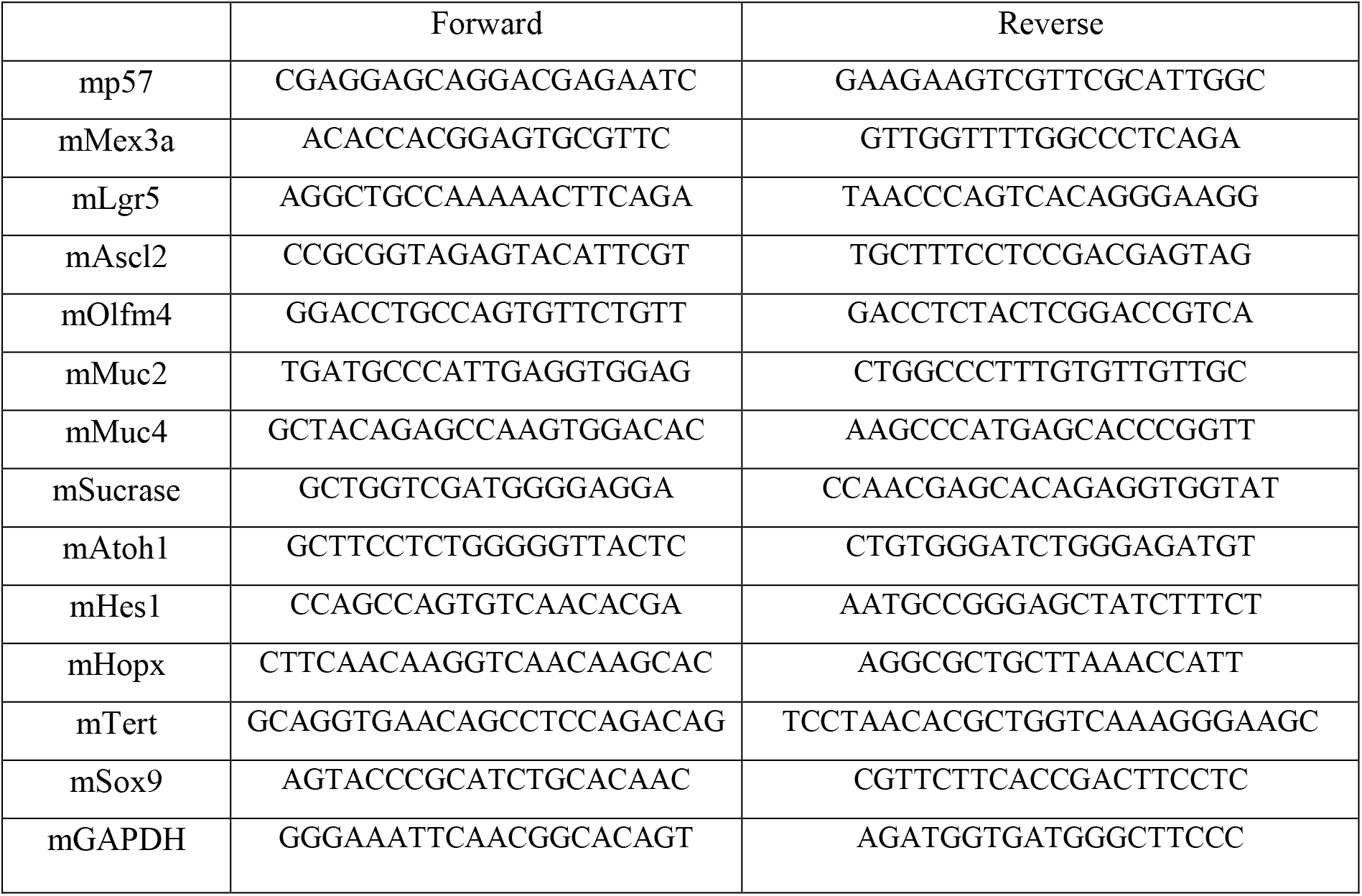

### Co-immunoprecipitation

Cells were scrapped and lysed in IP buffer (HEPES 50 mM pH 7.5, NaCl 150 mM, EDTA 1 mM, EGTA 2.5 mM, NP-40 1%, Tween-20 0.1%, glycerol 10%, supplemented with, dithiothreitol 1 mM, phosphatase inhibitors (β-glycerophosphate, NaF and sodium orthovanadate at 10 mM each) and protease inhibitors (Aprotinin, Bestatin, Leupeptin and Pepstatin A, each at 10 µg/mL). Lysates were sonicated for 10 s and centrifuged at 12500 g for 5 min at 4°C. Supernatants were collected and the protein concentration was determined by Bradford assay. The lysates (1 mg for SW480, 200 µg for HEK293) were incubated with 3 µg of the indicated antibodies and 12 µL protein sepharose beads (IPA300, Repligen) at 4°C for 4 h. Beads were washed 4 times with 750 µL of IP buffer and resuspended in 12 µL of Laemmli buffer, boiled for 3 min at 96°C and analyzed by immunoblotting as described below.

### DNA pull-down assays

For DNA pull-down, 80-mer oligonucleotides biotinylated either on the 5’ or 3’ end containing the E-box of the Lgr5 promoter or a mutated E-box (see table) were annealed with the corresponding antisense sequence and bound to streptavidin agarose beads (S1638, Millipore). Beads were washed 3 times, blocked for 3 h with salmon sperm DNA at a final concentration of 4 µg/µl and washed again 3 times with IP buffer.

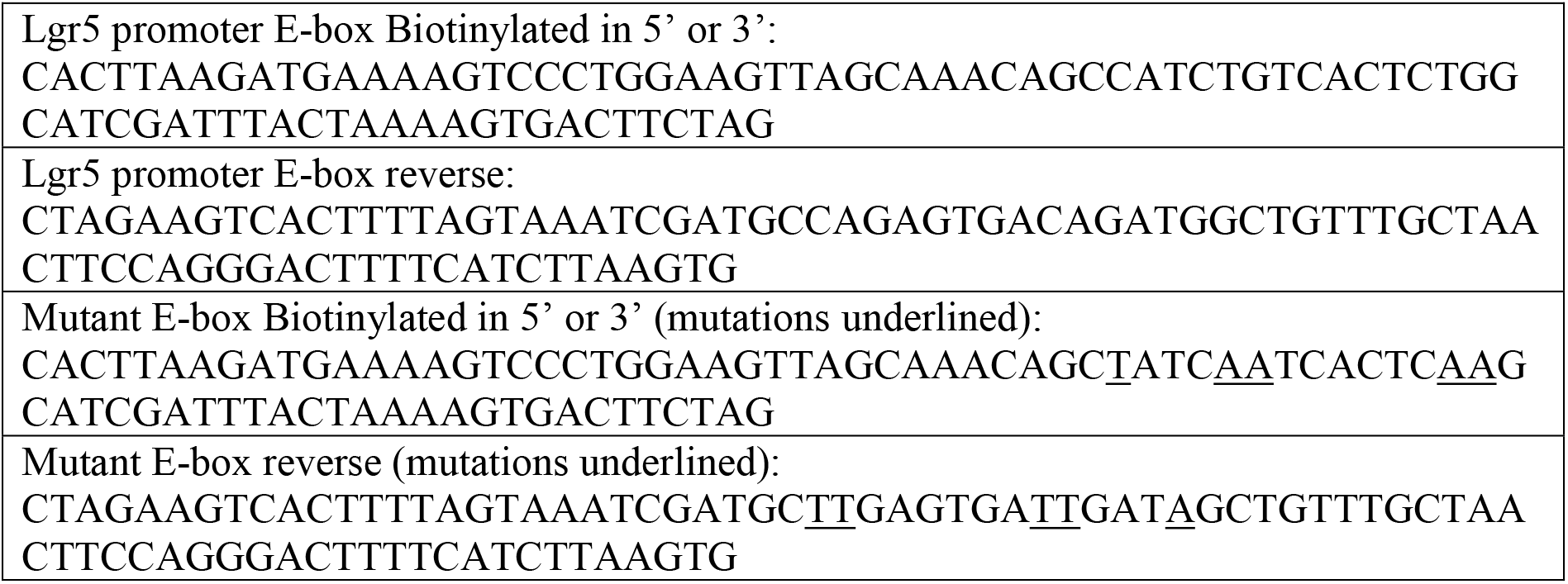

HEK 293 cells transfected with the indicated plasmids by the calcium phosphate method for 24 h in P100 plates and lysed in 800 µl of IP buffer as described above. Each pull-down consisted of 500 µl of IP buffer, 12 µl of beads slurry and 80 µl of protein lysate. These mixes were incubated under gentle agitation for 4 h at 4°C, rinsed 3 times with IP buffer and resolved on SDS-PAGE. After transfer on polyvinylidene difluoride membrane (PVDF) membranes, pull-downs were analyzed by probing with the indicated antibodies as described below.

### Immunoblotting

Lysates and immunoprecipitates were prepared as described above. Proteins were resolved on 8 to 12% SDS-PAGE gels, depending on protein size, and transferred to PVDF (Immobilon-P, Millipore). Membranes were blocked for 1 h in PBS-T (PBS 0.1% Tween 20), 5% non-fat dry milk and probed with primary antibodies overnight at 4°C under agitation. After three washes in PBS-T, membranes were probed with HRP-conjugated secondary antibodies at 1/10000 dilution. Signals were visualized with chemiluminescence detection reagent (Sigma, Bio-Rad) and autoradiography films (Blue Devil) or on a Fusion Solo S (Vilber) digital acquisition system.

### GFP transcription reporter assays

HEK293 cells were seeded in 24-well plates (45000 cells/well) and co-transfected with 100 ng of reporter plasmid (pZsGreen DA [without promoter, negative control], pZsGreen pGAPDH [positive control], pZsGreen pPhox2a [Ascl1 target, negative control], pZsGreen pSox9 or pZsGreen pLgr5 [Ascl2 targets]), 100 ng of pcDNA3.1+ Hygro Ascl2 plasmid and increasing amounts (0, 100 or 200 ng) of pcDNA3.1+ Hygro p57^WT^ or pcDNA3.1+ Hygro p57^CK-^ plasmid. Medium was changed 24 h after transfection and 24 h later, 9 images per well were captured in phase contrast and fluorescence with an Incucyte FLR automated microscope (Essen Bioscience) equipped with a 20X objective. The number of fluorescent cells (Object confluence) was quantified on each image using the Incucyte Software (v2011A) and values were normalized by the number fluorescent cells obtained with the pGAPDH reporter plasmid in the same condition.

### Chromatin immunoprecipitation

SW480 cells were grown to confluence. Medium was replaced with PBS and cells were fixed with 1% formaldehyde for 10 min under agitation. Crosslink was stopped by adding glycine at a final concentration of 125 mM for 5 min. Cells were rinsed with cold PBS and scrapped with PBS containing protease inhibitors (Aprotinin, Bestatin, Leupeptin and Pepstatin A at 10 µg/mL each). After centrifugation for 10 min at 3000 rpm, cell pellets were resuspended and lysed on ice for 30 min using 200 µL of lysis buffer per 1.10^6^ cells (10 mM Tris-HCl pH 8, 0.25% Triton-X100, 10 mM EDTA, 0.5 mM EGTA, 10 mM sodium butyrate, supplemented with 20 mM β-glycerophosphate, 10 µM sodium orthovanadate and protease inhibitors at 10 µg/mL). Cells were pipetted 20 times in a Dounce homogenizer on ice. The lysate was centrifuged for 5 min at 3000 rpm and the pellet containing DNA and cross-linked proteins was resuspended in 300 µL sonication buffer per 3.10^6^ cells (10 mM Tris-HCl pH 8, 100 mM NaCl, 1 mM EDTA, 0.5 mM EGTA, 10 mM Sodium butyrate, supplemented with 20 mM β-glycerophosphate, 10 µM sodium orthovanadate, 1% SDS and protease inhibitors at 10 µg/mL) and incubated for 10 min on ice. Chromatin was sheared by sonication (16 cycles of 30 s On/20 s Off, 40% amplitude using a Vibra-cell VCX130 Sonicator [Sonics]) on ice. Samples were centrifuged for 10 min at 1400 rpm to eliminate SDS, and the supernatants containing DNA were transferred to new tubes. Chromatin shearing was verified on agarose gel and DNA concentration was determined with a Nanodrop spectrophotometer. For each ChIP, 200 µg of chromatin were used, and 10% of the volume was kept as control input. The volume of IP was completed to 800 µL with sonication buffer and converted to RIPA buffer by adding 80 µL of 10% Triton-X100, 23 µL of 5 M NaCl and 8 µL of sodium deoxycholate (DOC). For each IP, 4 µg of antibodies and 20 µL of Magna Chip Protein A/G magnetic beads (Millipore) were added and samples were incubated overnight on a rotating wheel at 4°C. Samples were washed 3 times with low salt buffer (10 mM Tris-HCl pH 8, 0.1% Triton X-100, 0.1% SDS, 0.1% DOC, 140 mM NaCl, 1 mM EDTA, 0.5 mM EGTA, 10 mM Sodium butyrate, supplemented with 20 mM β-glycerophosphate and 100 µM sodium orthovanadate), three times with high-salt buffer (same as low salt but with 500 mM NaCl), twice with LiCl buffer (0.25 M LiCl, 1% NP-40, 1% DOC, 10 mM Tris-HCl pH 8, 1 mM EDTA, 10 mM Sodium butyrate and 100 µM sodium orthovanadate) and twice with TE buffer (10 mM Tris-HCl, 1 mM EDTA, pH 8). DNA purification and elution was realized with the IPure v2 kit (Diagenode) according to the manufacturer’s protocol. qPCRs were performed with 10 ng of DNA per reaction as described above using the following primers.

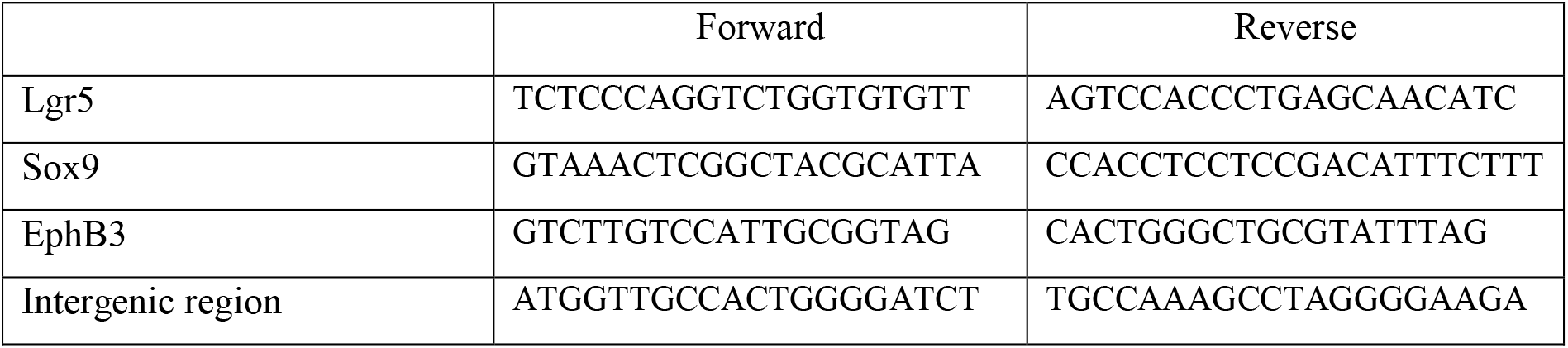

### Proximity Ligation Assay

PLAs were performed using the Duolink in situ fluorescence technology kit (Sigma) according to the manufacturer’s instructions. Briefly, SW480 were seeded on glass coverslips at 25000 cells/cm^2^ and grown for 2 days. Cells were fixed with 4% PFA for 20 min at 37°C and permeabilized with PBS 0.2% Triton-X100 for 3 min. Coverslips were blocked at 37°C with Duolink blocking solution and incubated with the indicated combinations of primary antibodies for 1 h at 37°C. Coverslips were washed twice in Duolink wash buffer A and incubated with PLA probes, anti-Rabbit PLUS (DUO92002) and anti-Mouse MINUS (DUO92004) for 1 h at 37°C. Duolink detection reagents were used for signal detection. Following the PLA procedure, cells were stained with Phalloidin A488 at 1/500 for 30 min at 37°C and DNA was stained with Hoechst 33342 (0.1 µg/mL). Images were acquired on a Nikon 90i Eclipse microscope using a DS-Qi2 HQ camera. The following antibodies were used in PLA experiments: rabbit anti-p57 (Sigma, P0357, 1/500), Ascl2 (Sigma, SAB1305026, 1/100), HDAC7 (Sigma, SAB4502143, 1/100) and mouse anti-mSin3a (G-11, sc-5299, 1/100), HDAC7 (A-7, sc-74563, 1/100), Ascl2 (Millipore, 8F1, MAB4417, 1/100).

### Quantification and statistical analysis

In all figures, n represents the number of independent experiments. Flow cytometry and cell sorting experiments were performed with blind p57 genotype. Statistical analyzes were performed using GraphPad Prism 6.0. Differences between three groups or more were evaluated by ANOVA test followed by Bonferroni multiple comparison test. Differences between two groups were evaluated by unpaired t-test with the post-hoc test of Mann-Whitney. Data are presented as ± s.e.m. Symbols used are: ns: p > 0.05; *: p ≤ 0.05; **: p ≤ 0.01; ***: p ≤ 0.001; ****: p ≤ 0.0001.

### Transcriptomic analyses

Ten RNA-seq samples were sequenced using Illumina HiSeq (paired-end, 2×150 bp reads) at Genewiz. The quality of each raw sequencing file (fastq) was verified with FastQC (https://www.bioinformatics.babraham.ac.uk/projects/fastqc/). For all files adapters were trimmed with Trim Galore (v-0.6.5, https://www.bioinformatics.babraham.ac.uk/projects/trim_galore) and clean fastq were aligned to the reference mouse genome (mm10) using STAR aligner (STAR_2.6.1d) [73]. FPKM were computed using cufflinks (v2.2.1) [74] with the annotation from Ensembl database (gtf GRCm38.91). Raw read count per sample was computed using HT-seq count (V1.0) [75] on that same annotation database. Then the raw count table was cleaned, and only genes with a total higher than 50 reads aligned over all samples were kept. The remaining raw read count were normalized using the RLE methods and differential analysis was applied per pairs of conditions using DESeq2 (DESeq2 package version: 1.22.2) [76], available as an R package in bioconductor (www.biocondutor.org), to generate the log fold change (log2FC) values and adjusted p-values (Benjamini and Hochberg method) (Table S2). Heatmaps were drawn using R, on FPKM values.

RNA-Seq data have been deposited on the GEO database under accession number GSE179664.

## Data Availability

RNA-Seq data have been deposited on the GEO database under accession number GSE179664.

## Acknowledgements

We thank Dr Philippe Jay and Julie Nguyen (IGF, Montpellier) for expert advice on FACS sorting of ICSs, Dr Ludivine Drougat (CBI, Toulouse) for useful discussion for RNA-Seq data analysis and interpretation, and Dr Fabrice Escaffit (CBI, Toulouse) for advice on RT-qPCR. We are grateful to Dr. Eric Verdin (UCSF, San Francisco) and Dr. Robert Eisenman (Fred Hutchinson Cancer Research Center, Seattle) for providing reagents. The authors thank the personnel of the ABC animal facility.

## Author contributions

JC, AN, APr, MA, ND, TJ, CD and AB designed experiments. JC, AN, APr, APi, ND, CC, and AB performed experiments. JC, AN, MA, APr, APi, ND and AB analyzed the data. JC and AB wrote the paper with contributions from all authors. AB acquired the funds and supervised the work.

## Figure Legends

**Figure S1:**
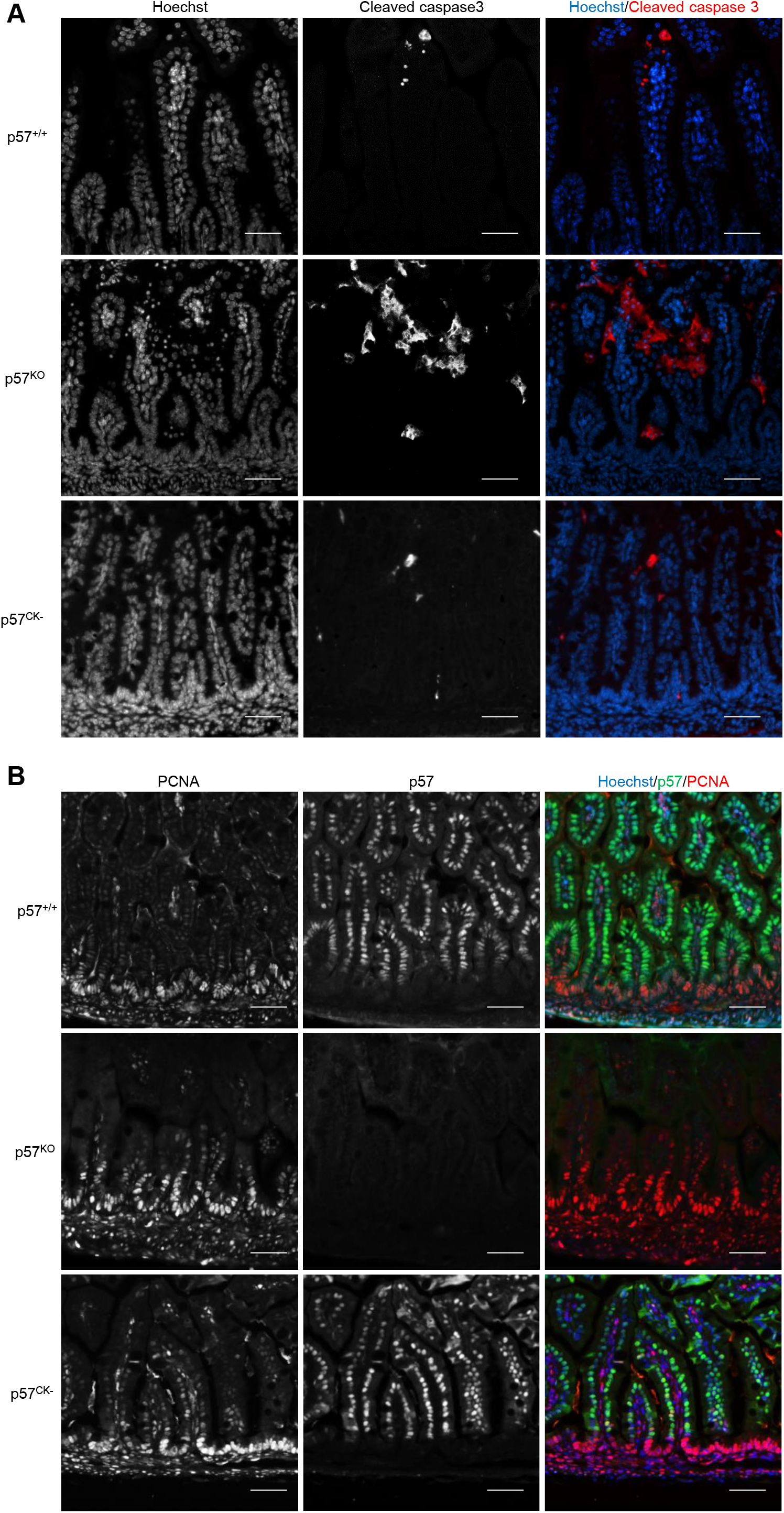
Increased apoptosis in villi and increased proliferation in crypts of p57^KO^ intestines. Intestine sections of E18.5 p57^+/+^, p57^KO^ and p57^CK-^ embryos were stained for cleaved caspase 3 (A) or p57 and PCNA (B) to assess apoptosis and proliferation, respectively. DNA was stained with Hoechst 33342. p57^KO^ intestine exhibit increased apoptosis in villi and increased proliferation in intervillous domains. This phenotype was never observed in p57^CK-^ mutant mice. Scale bars: 50 µm.

**Figure S2:**
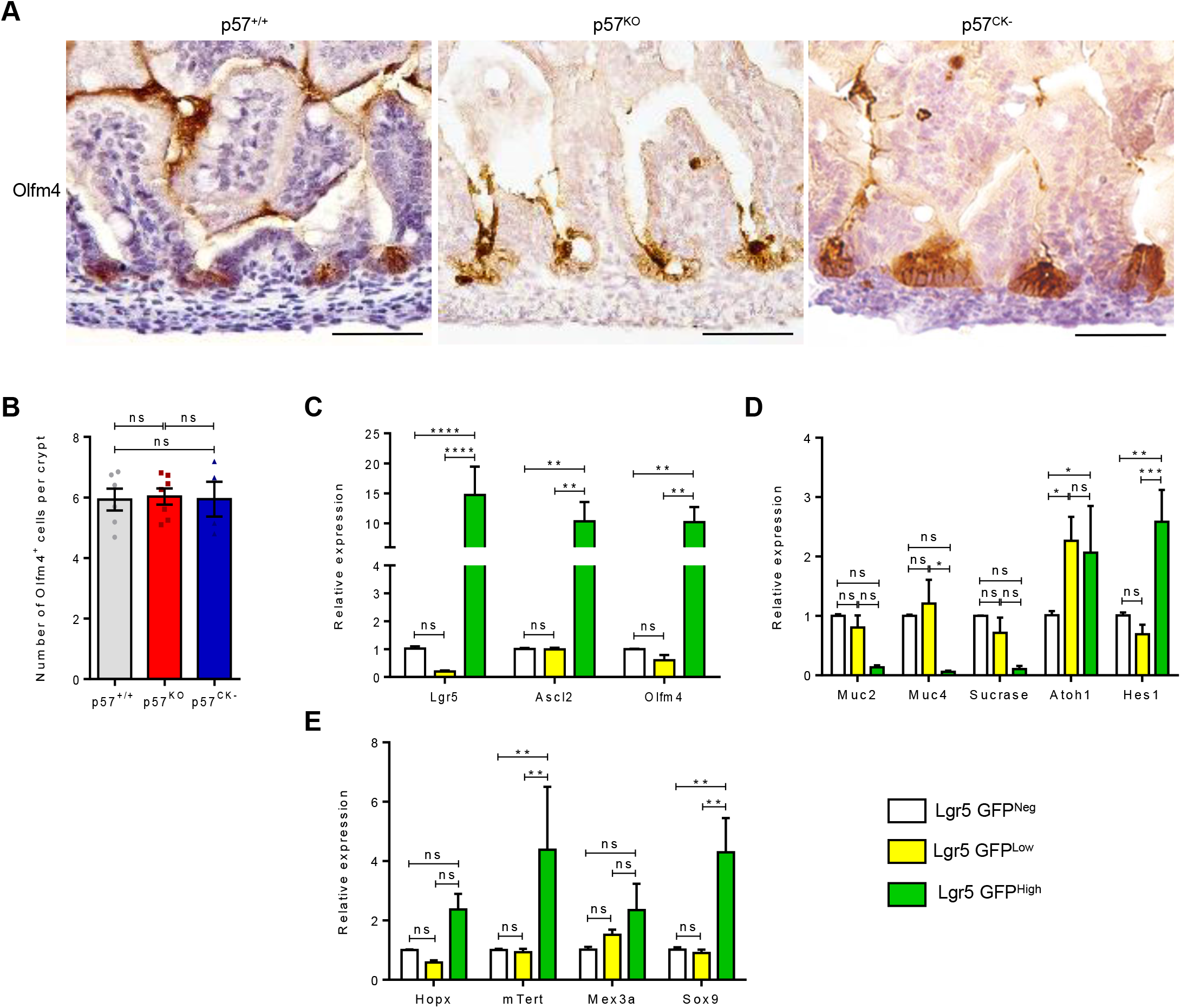
Characterization of CBCs in p57 mutant mice. (A) Immunohistochemistry (brown) for the CBC marker Olfm4 in intestine sections from p57^+/+^, p57^KO^ and p57^CK-^ E18.5 embryos. Scale bars: 50 µm. (B) Quantification of the number of Olfm4^+^ cells per intervillous domain in p57^+/+^ (n=6), p57^KO^ (n=7) and p57^CK-^ (n=4) intestines as described in A. (C-E) Verification of the molecular identity of CBCs by RT-qPCR on sorted intestinal epithelial cells. Dissociated crypts from Lgr5^+/eGFP-CreERT^ intestines were sorted based on GFP expression and gene expression of known CBC (C), differentiation (D) and reserve ISC (E) markers was evaluated in Lgr5 GFP^Neg^, Lgr5 GFP^Low^ and Lgr5 GFP^High^ populations (n=3).

**Figure S3:**
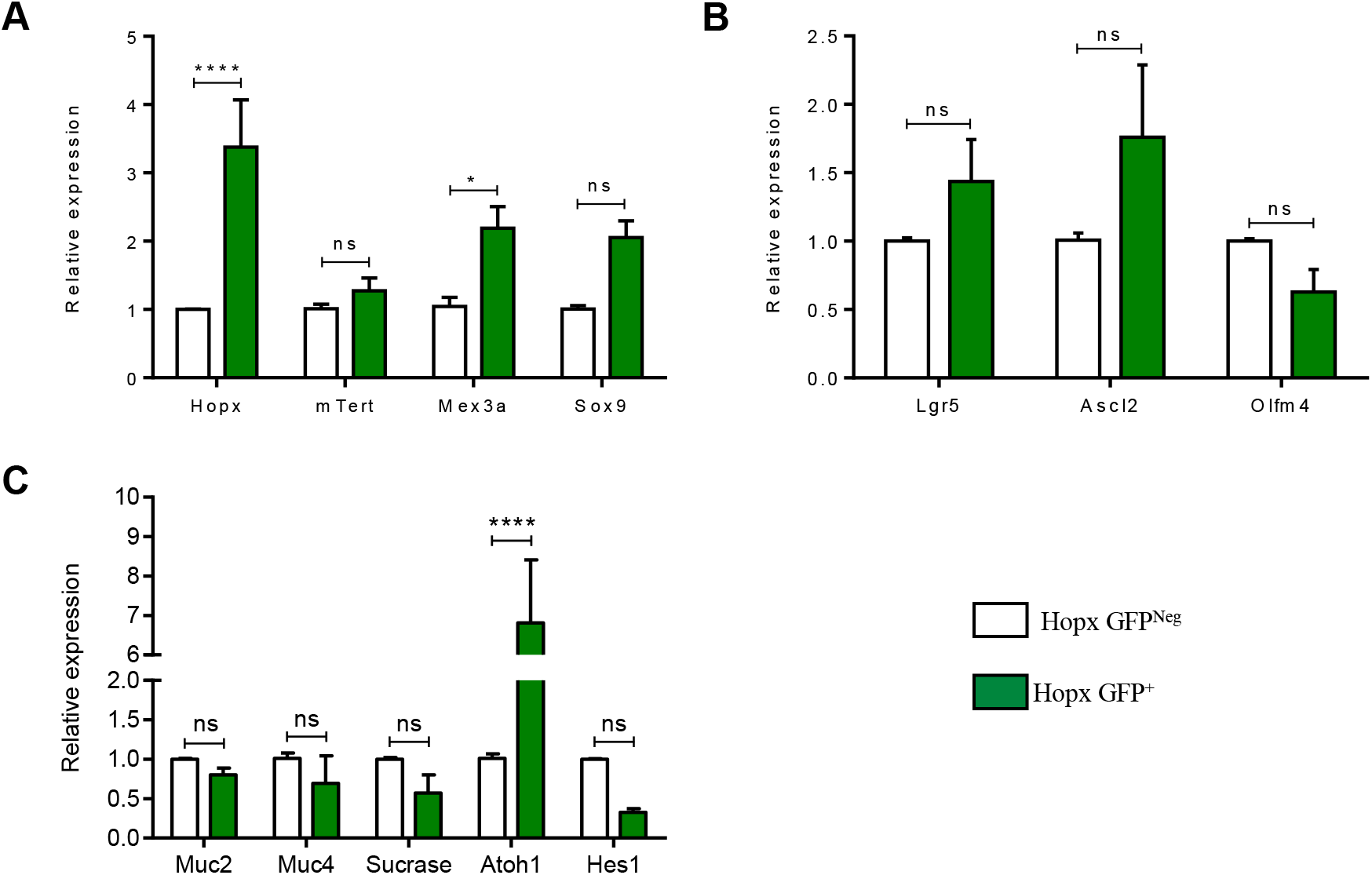
Characterization of Hopx^+^ ISCs in p57 mutant mice. (A-C) The molecular identity of Hopx^+^ ISCs was verified by RT-qPCR on sorted intestinal epithelial cells. Dissociated crypts from Hopx^3Xflag-eGFP^ intestines were sorted based on GFP expression and gene expression of known markers of reserve ISCs (A), CBCs (B) and differentiation (C) was evaluated in Hopx-GFP^Neg^ and Hopx GFP^+^ populations (n=3).

**Figure S4:**
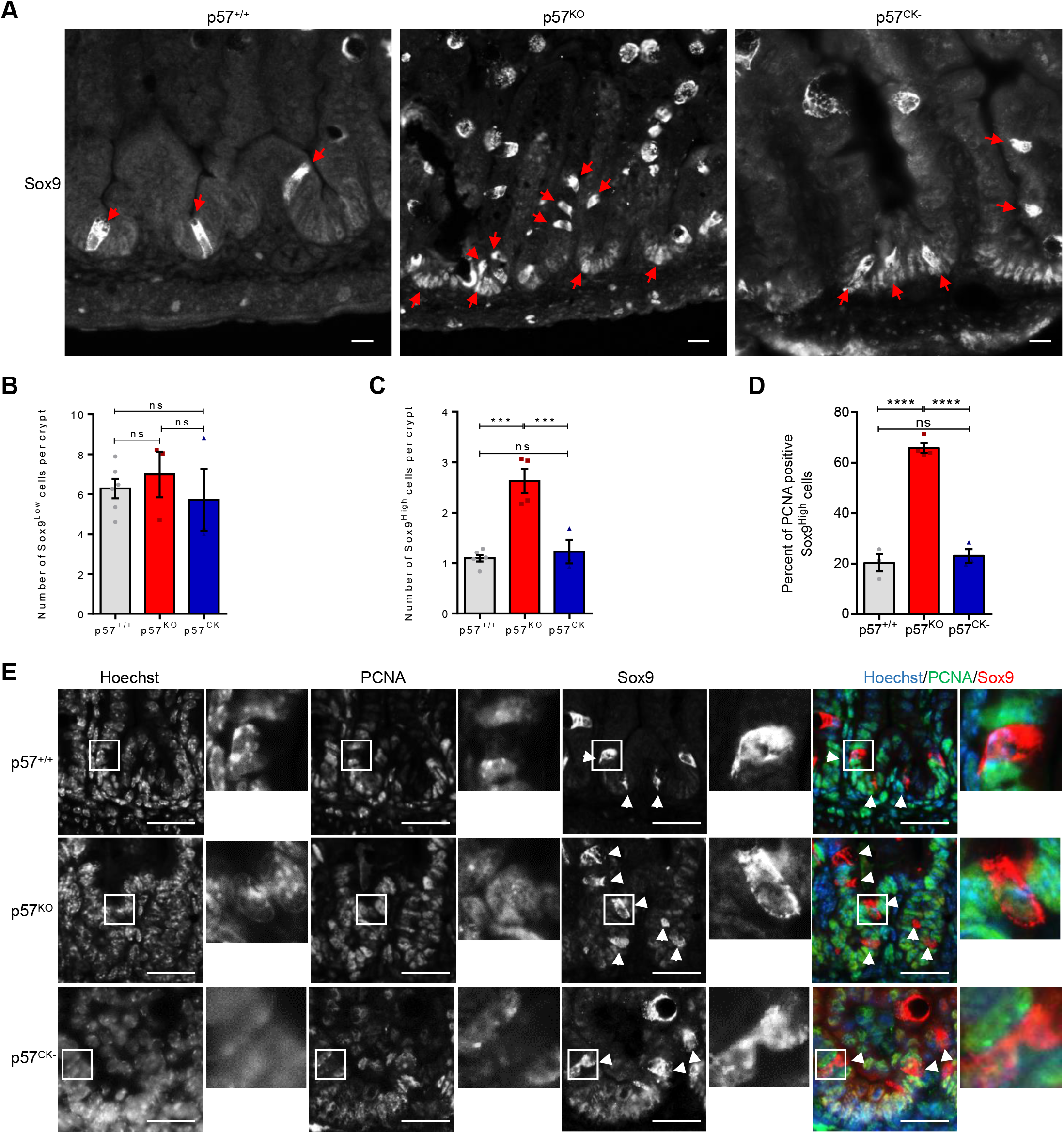
Amplification of proliferative of Sox9^High^ cells in p57^KO^ intestines. (A) Intestine sections from p57^+/+^, p57^KO^ and p57^CK-^ E18.5 embryos were immunostained for Sox9. CBCs express low levels of Sox9 while reserve ISCs express high levels of Sox9 (arrowheads). Goblet cells also exhibit high Sox9 expression but can be distinguished from reserve ISCs by their distinct morphology. Scale bars: 50 µm. (B-C) Quantification of Sox9^Low^ (B) and Sox9^High^ (C) cells per intervillous domain in p57^+/+^ (n=6), p57^KO^ (n=3) and p57^CK-^ (n=3) E18.5 intestines as described in A. (D) Percentage of proliferating Sox9^High^ cells in intestinal sections immunostained for Sox9 and PCNA from p57^+/+^ (n=3), p57^KO^ (n=3) and p57^CK-^ (n=3) E18.5 embryos, as shown in E. (E) Representative images of Sox9 and PCNA immunostaining of E18.5 intestine sections from p57^+/+^, p57^KO^ and p57^CK-^ E18.5 embryos. Sox9^High^ cells are indicated by arrowheads. DNA was stained with Hoechst 33342. Scale bars: 20 µm.

**Figure S5:**
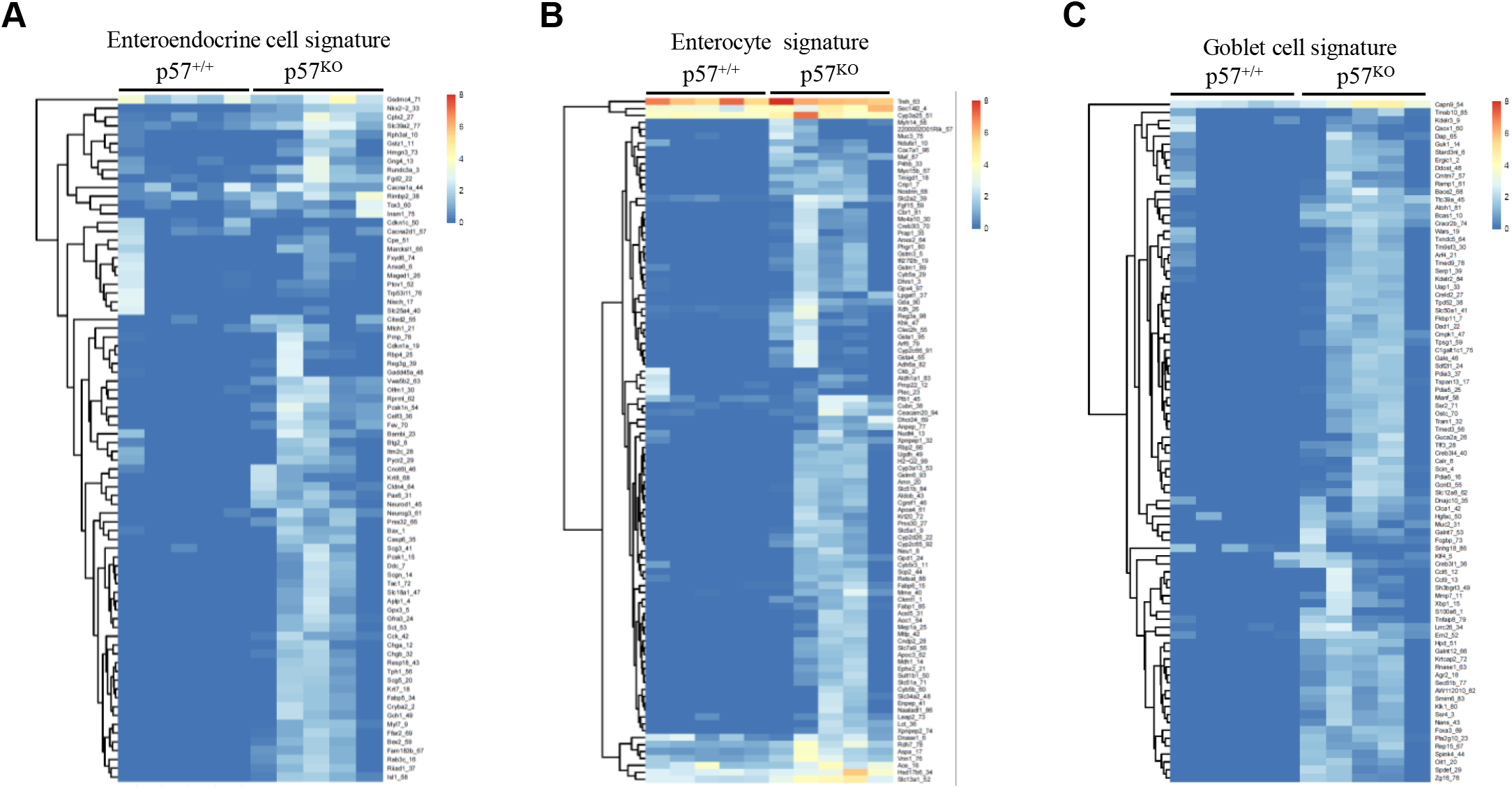
Alteration of gene expression signatures in Hopx^+^ ISCs of p57^KO^ mice. (A-C) Heatmaps of Hopx-GFP^+^ ISC RNA-Seq analyses from p57^+/+^ (n=5) and p57^KO^ (n=5) mice for previously published gene expression signatures of specific intestinal cell populations: Enteroendocrine cells (A), Enterocytes (B), and Goblet cells (C).

**Figure S6:**
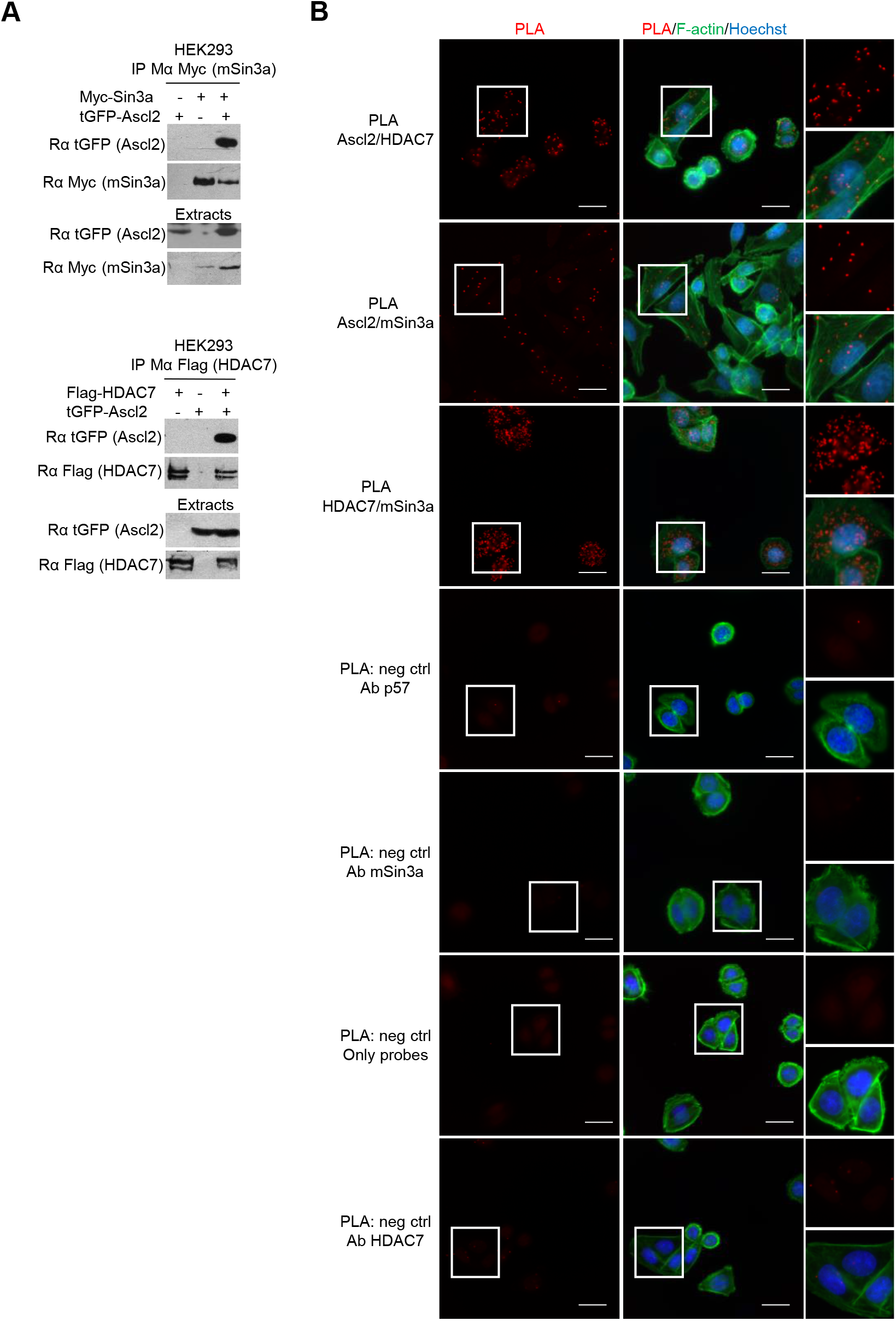
Validation of the interaction partners of the co-repressor complex. (A) mSin3a or HDAC7 were immunoprecipitated with Myc or Flag antibodies, respectively, from HEK293 cells transfected with tGFP-Ascl2 and/or Myc-mSin3a and/or Flag-HDAC7. The presence of Ascl2 was assessed by immunoblotting with the indicated antibodies (n=3). Respective levels of transfected proteins in the extracts are shown. (B) PLA using Ascl2, mSin3a and/or HDAC7 antibodies on endogenous proteins in SW480 cells. F-actin was stained with phalloïdin and DNA with Hoechst 33342. Scale bars: 50 µm. Representative images of three independent experiments.

**Figure S7:**
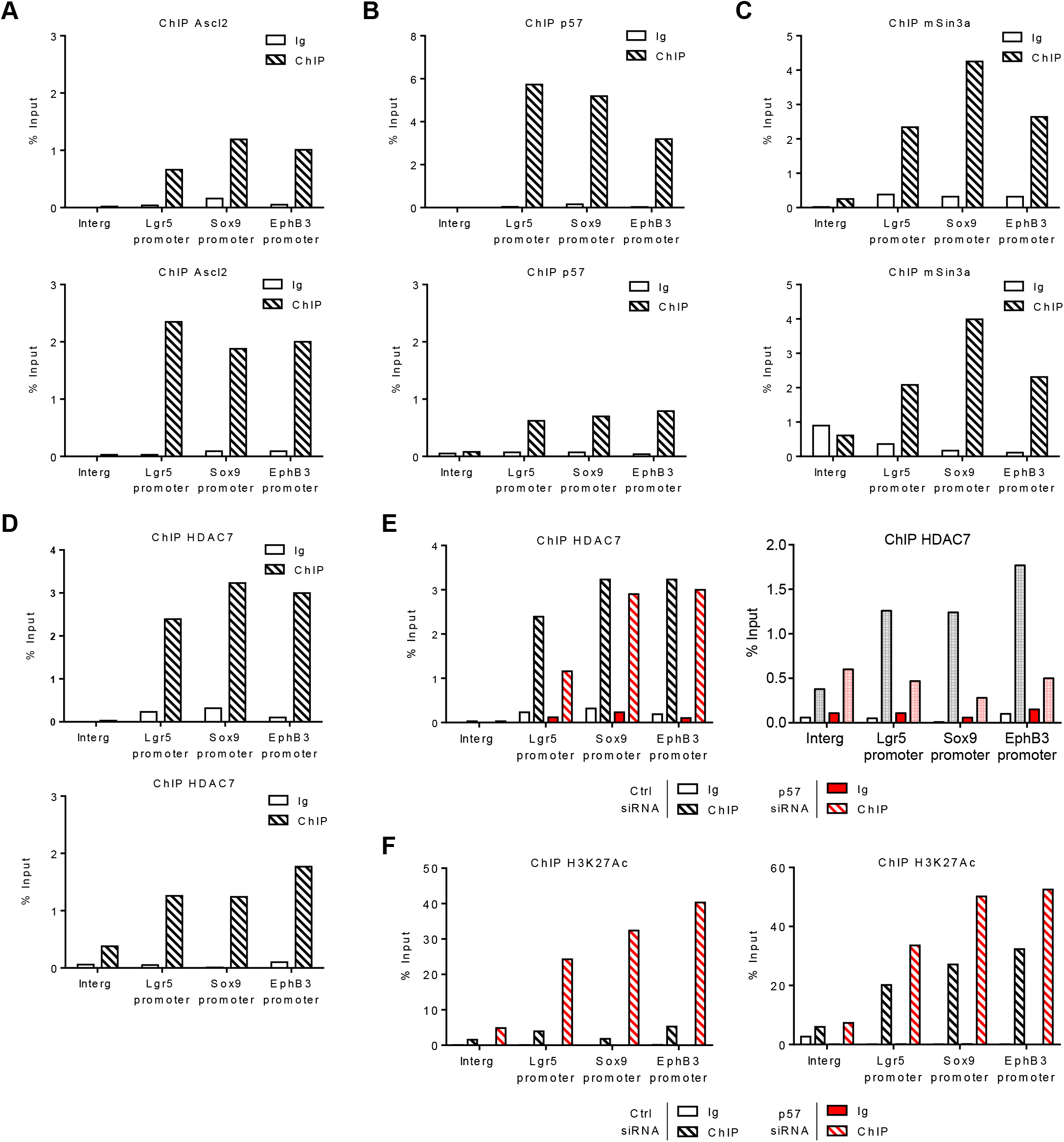
p57 participate in the recruitment of a co-repressor complex on Ascl2 target promoters. (A-F) Graphs show the results of two assays from three independent experiments, the third one is displayed in Figure 8. (A-D) ChIPs were performed with antibodies against Ascl2 (A), p57 (B), mSin3a (C), HDAC7 (D) or IgG as negative control to determine the presence of the co-repressor complex on Ascl2 target promoters. (E-F) ChIPs with anti HDAC7 (K) or H3K27Ac (L) antibodies or control IgG in SW480 cells transfected for 48 h with control or p57 siRNA followed by qPCR with primers specific for the Lgr5, Sox9 and EphB3 promoters or for an unrelated intergenic region as negative control.

**Table S1:** Intestine length in mm from p57^+/+^, p57^KO^ and p57^CK-^ E18.5 embryos.

| p57<sup>+/+</sup> | p57<sup>KO</sup> | p57<sup>CK-</sup> |
| --- | --- | --- |
| 121 | 34 | 72 |
| 123 | 6 | 84 |
| 112 | 22 | 103 |
| 114 | 105 | 77 |
| 119 | 92 | 108 |
| 122 | 107 | 87 |
| 103 | 110 | 79 |
| 111 | 100 | 114 |
| 94 | 110 | 94 |
| 95 | 100 | 50 |
| 99 | 90 | 132 |
| 93 | 61 | 153 |
| 98 | 77 | 124 |
| 102 | 58 | 110 |
| 108 | 68 | 130 |
| 99 | 88 | 112 |
| 75 | 87 | 122 |
| 72 | 90 | 104 |
| 86 | 91 | 91 |
| 80 | 20 | 113 |
| 92 | 61 | 99 |
| 83 | 64 | 103 |
| 95 | 75 | 91 |
| 91 | 90 | 106 |
| 103 | 98 | 117 |
| 105 | 60 |  |
| 96 | 81 |  |
| 99 | 89 |  |
| 110 | 70 |  |
| 94 |  |  |
| 95 |  |  |
| 107 |  |  |
| 83 |  |  |
| 100 |  |  |
| 107 |  |  |
| 83 |  |  |
| 100 |  |  |
| 92 |  |  |
| 86 |  |  |
| 89 |  |  |
| 86 |  |  |
| 87 |  |  |
| 67 |  |  |
| 65 |  |  |
| 92 |  |  |
| 104 |  |  |
| 97 |  |  |
| 79 |  |  |
| 76 |  |  |
| 81 |  |  |
| 99 |  |  |
| 83 |  |  |
| 93 |  |  |
| 104 |  |  |
| 108 |  |  |
| 100 |  |  |
| 77 |  |  |
| 96 |  |  |
| 94 |  |  |
| 93 |  |  |

